# Spatially aligned random partition models on spatially resolved transcriptomics data

**DOI:** 10.1101/2025.04.16.649218

**Authors:** Yunshan Duan, Shuai Guo, Hao Yan, Wenyi Wang, Peter Mueller

## Abstract

We propose spatially aligned random partition (SARP) models for clustering multiple types of experimental units, incorporating dependence in a subvector of the cluster-specific parameters, e.g., a subvector of spatial information, as in the motivating application. The approach is developed for inference about co-localization of immune, stromal, and tumor cell sub-populations. The aim is to understand the recruitment of immune and stromal cell subtypes by tumor cells, formalized as spatial dependence of the corresponding homogeneous cell subpopulations. This is achieved by constructing Bayesian nonparametric random partition models for the different types of cells, with a hierarchically structured prior introducing the desired dependence. Specifically, we use Pitman-Yor priors and add dependence in the base measure for spatial features, while leaving the base measure corresponding to gene expression features a priori independent across different types of cells. Details of the model construction are designed to lead to a convenient MCMC algorithm for posterior inference. Simulation studies show favorable performance in identifying co-localization between types of cells. We apply the proposed approach with colorectal cancer (CRC) data and discover subtypes of immune and stromal cells that are spatially aligned with specific tumor regions.

## 1 Introduction

Current literature on Bayesian inference for random partitions includes extensive work on joint models for multiple random partitions allowing for dependence. However, there is no principled approach for inference about specific clusters in a partition of one type of experimental units relating to clusters in a random partition of another type of experimental units. We require such inference for a motivating application to spatial transcriptomics data, to answer questions about specific subtypes of tumor cells recruiting specific immune cell subtypes. To address this question, we develop a Bayesian nonparametric (BNP) inference approach for spatially aligned random partitions across multiple sets of experimental units. That is, the proposed inference implements model-based clustering on separate sets of units, introducing dependence in a subvector of the cluster specific parameters corresponding to spatial location (or any other shared feature sub-space – for simplicity, without loss of generality, we will throughout assume spatial dependence). In the motivating application, the experimental units are immune, stromal, and tumor cells from tumor tissue, with the inference question being how subtypes of these cells co-localize with one another. We integrate spatial transcriptomics (ST) and single-cell RNA sequencing (SC; scRNA-seq) data to obtain both gene expression profiles and spatial coordinates for individual cells. Motivated by the observation that immune and stromal cells often cluster around specific tumor regions, we aim to characterize the spatial co-localization patterns among subpopulations of these three cell types. In our framework, the spatial alignment of cell subtypes is formalized as dependence in random partitions of the spatial subvector, while gene expression remains independent. In the motivating case study such inference is implemented to better understand the tumor microenvironment and the immunological interactions that influence tumor progression.

Several methods have been proposed for spatially informed clustering in spatial transcriptomics data, including approaches such as BayesSpace [1], SpaGCN [2], Starfysh [3] and SpaRTaCo [4]. For example, SpaRTaCo jointly clusters genes and spatial tissue regions by inferring a latent block structure from spatial gene expression data, enabling co-clustering of genes and spots for biological interpretation. However, these methods focus on clustering within a single set of spatial units and do not support inference on the alignment or interaction between distinct partitions across multiple cell types. In contrast, our framework enables such cross-partition inference by inducing dependence across spatial cluster structures associated with different cell types.

We implement the desired inference under a Bayesian paradigm. Bayesian inference on random partitions is often implemented by way of setting up a mixture model 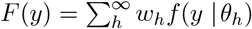. For statistical inference it is convenient to write *F*(*y*) as a mixture of the kernel *f*(*y* | *θ*) with respect to a discrete mixing measure 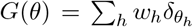. Here *δ*_*θ*_ is a point mass at *θ*. Under a Bayesian approach the model construction is completed with a prior probability model for the random mixing measure *G*. Prior models for random probability measures are known as non-parametric Bayesian (BNP) models [5]. The most widely used BNP prior, specifically for model-based clustering is the Dirichlet process (DP) prior [5, Chapter 4]. However, the law of the random partition induced by the DP, known as the Chinese restaurant process (CRP), is controlled by only a single parameter [6]. Several alternative models generalize DP mixture models to allow for more flexibility in clustering, using, for example, the Pitman-Yor process (PYP) [7], and the normalized inverse Gaussian as special cases [8].

Cancer tissues consist of diverse populations of epithelial, stromal, and immune cells, each encompassing multiple distinct cell types and characteristics. Gaining insights into the specific roles of these cellular components and their interactions is essential for furthering our understanding of cancer biology. A vast majority of solid tumors originate from epithelial cells, therefore, throughout the paper, epithelial is used as a proxy for the tumor cell population. To better understand the interaction of different cell types, we set up a random partition of tumor, immune, and stromal cells, allowing for spatial correlation of different subtypes. Specifically, we represent cell populations composed of homogeneous cell subtypes by way of mixtures of kernels with respect to mixing measures *G*_*j*_, *j* = 1, 2, 3, for tumor, stromal and immune cells, respectively. In this context, we represent the recruitment of immune and stromal cell subtypes by tumor cells as spatial dependence. In the mixture model this becomes spatial dependence of the locations of (some) immune and stromal cell subpopulations with the locations of tumor cell subpopulations. The desired dependence is formalized as a dependent prior *p*(*G*_1_, *G*_2_, *G*_3_) across the mixtures. Many dependent priors for a family 𝒢 = {*G*_1_,…, *G*_*J*_} of random probability measures have been studied in the BNP literature over the past 25 years. Some of the earliest constructions introduce the desired dependence by way of shared hyperparameters [9, 10]. Another early construction is based on additive decomposition models using a mixture of common versus idiosyncratic parts based on DPs [11]. Variations to ensure DP marginal distribution are proposed in Kolossiatis et al. [12], and extensions to completely random measures (CRM) [13] are introduced in Lijoi et al. [14]. Other types of widely used dependent models include the hierarchical DP (HDP) [15], the nested DP (NDP) [16] and their variations. Another widely used construction for dependent random probability measures is introduced in the seminal paper by MacEachern [17] as the dependent DP (DDP) model for problems when the random probability measures are indexed by covariates as 𝒢 = {*G*_*x*_; *x* ∈ *X*}. A discussion on the properties of the various dependent priors constructions has been included in [18].

However, the motivating application in the upcoming discussion requires more structure than what is naturally accommodated in these approaches. For example, there is an asymmetry in the desired inference. Also dependence is restricted to a spatial subvector of linked subpopulations of the immune, stromal, and tumor cells, but gene expression profiles should remain a priori independent. None of the earlier mentioned constructions favors such structure as we need it here for a meaningful biological interpretation. This motivates us to propose spatially aligned random partition models that introduce the co-localization of clusters with the desired structure. The proposed framework approaches the problem as an inference problem with dependent partitions, building on the rich literature of dependent random partition models that we briefly reviewed before. However, other approaches are possible. One alternative and very natural approach is to build on an equally rich literature of point processes. This direction is explored, for example, in Osher et al. [19].

We propose spatially aligned random partition (SARP) models to identify clusters of multiple cell types within the same spatial context. The random partition (or clustering) of cells is informed by both gene expression profiles—summarized using principal components (PCAs)—and spatial coordinates, enabling us to identify biologically relevant subpopulations that are coherent in both molecular and spatial domains. The desired structure in constructing dependent random partitions for this application includes that (1) aligned clusters should exhibit similar but not identical spatial locations; (2) the dependence structure should allow for one type of cells (e.g., tumor cells) to serve as reference with clusters of other types of cells (e.g., immune cells and stromal cells) to be located around reference cell clusters; and (3) alignment should allow multiple clusters of the other types to be spatially aligned with one subtype of the reference type; (4) the alignment is introduced only for a subvector of the features – in our case, the spatial coordinates. The desired structure is achieved by adding dependence in the base measure of Pitman-Yor priors. The construction allows for a natural and efficient posterior simulation algorithm. We apply the proposed SARP model for inference with colorectal cancer (CRC) data.

The rest of the paper is organized as follows. Section 2 describes the motivating application of studying cell interaction in the tumor micro-environment and the data source. We describe the proposed pipeline and data pre-processing. In Section 3.1 we introduce the proposed SARP model, first for the case of two sets of experimental units. Section 3.2 discusses marginal and asymptotic properties of the model. Section 3.3 extends the model to more than two sets of units, and Section 3.4 introduces the posterior inference algorithm. In Section 4, we summarize simulation studies to validate the proposed inference approach. In Section 5, we apply the proposed method for inference with colorectal cancer (CRC) data. The results find immune and stromal cell subtypes spatially aligned with tumor subtypes. R code including implementation of the proposed method and all results in the paper is available in Github repository https://github.com/YunshanDYS/SARP.

## 2 Case Study

CRC is the second most common cause of cancer death in the United States with approximately 153,020 new cases in 2023 [20]. It is well-established that immune cells within the tumor microenvironment (TME) profoundly influence tumor development, progression, and treatment responses in colorectal cancer (CRC) patients. Histological analyses have observed that immune and stromal cells often position themselves around specific tumor regions, suggesting intricate interactions between these cell types. While such spatial relationships have been noted, comprehensive characterization of the precise spatial co-localization of immune, stromal, and tumor cell subpopulations in CRC using spatial transcriptomics (ST) data remains limited. Notably, recent studies have begun to address this gap: a study integrating spatial and single-cell transcriptomics revealed that CRC cells at the invasive front co-localize with macrophages, highlighting spatial intercellular communication between tumor cells and the immune microenvironment [21]. Another study demonstrated that the spatial organization and immune status of the tumor-stroma boundary are distinctive features of different CRC subtypes, influencing disease progression and therapeutic responses [22]. Building upon these insights, our objective is to utilize ST technology to systematically identify and characterize immune and stromal cell subpopulations that are spatially aligned with tumor cells. This approach aims to provide a more detailed understanding of cellular interactions within the CRC TME, potentially uncovering novel therapeutic targets or prognostic indicators.

We use data from two advanced RNA sequencing techniques, including single-cell RNA sequencing (SC) and spatial transcriptomics (ST) data. SC data is now widely used for analysis [23]. More recently, ST data has gained attention for enabling spatial inference of the TME by leveraging additional spatial location information [24, 25]. To assess the spatial interactions of specific cell types, we integrate information from both modalities during preprocessing to obtain spatially inferred single cell data for further analysis. The SC data is a publicly available CRC dataset from Lee et al. [26]. The ST sample is public domain data from the 10X website (https://www.10xgenomics.com/datasets/human-colorectal-cancer-11-mm-capture-area-ffpe-2-standard).

Figure 1 shows the workflow for the implemented pipeline. SC data provides gene expression profiles for each individual cell in the tumor tissue, while ST data includes both gene expression and spatial information for each spot on a tissue grid, representing the cumulative information of multiple cells within that spot. First, standard pre-processing and data cleaning is carried out for both, SC and ST data, using programs implemented in the *Seurat* package. For SC data, only genes that were detected in a minimum of three cells were retained. We excluded low-quality cells by setting thresholds for the number of expressed genes and detected Unique Molecular Identifier (UMI) counts: cells were retained if they expressed between 500 and 6,500 genes and had UMI counts ranging from 500 to 2,000,000. For ST data, genes with all 0s are removed. Additionally, we removed mitochondrial, microRNA, long non-coding, and predicted gene models to focus on high-confidence, protein-coding transcripts to avoid confounding signals from less informative or tissue-specific genes. Next, we use CellTrek [27] to spatially map the SC data leveraging the spatial annotation of ST data. The output from this step are single cells with gene expression and inferred spatial location on the tissue slide, i.e., spatially enriched SC data. Principle Component Analysis (PCA) is used for dimension reduction to *G* gene expression features. Uniform Manifold Approximation and Projection (UMAP) transformations are implemented for visualization (but not for analysis). Based on known markers for immune, stromal, and tumor cells, we split the spatially enriched SC data into three subsets by cell type. Figure 2 shows the spatially enriched SC data with cell type annotation in the CRC datasets. In the two subfigures, single cells are colored by the annotated cell types and plotted with respect to spatial locations and gene expression UMAP scores, respectively. Using these spatially enriched data, the inference goal is then the discovery of spatially aligned random partitions, aligned across tumor, immune and stromal cells.

**Figure 1:**
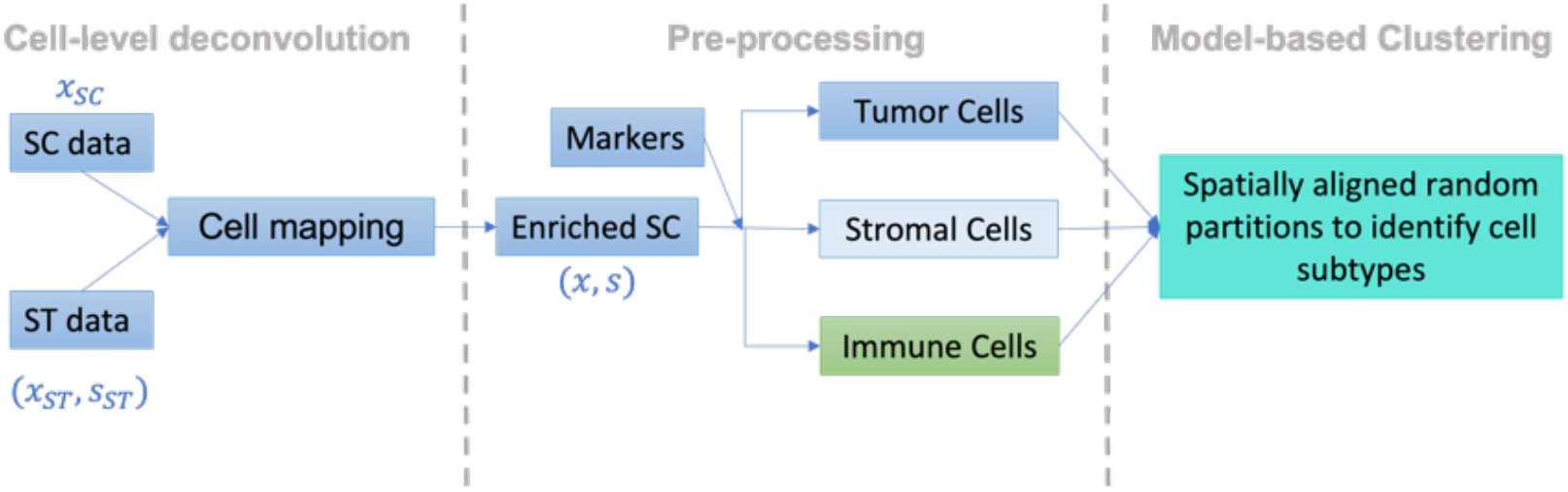
Workflow of the proposed method. Here *x*_SC_ denotes single cell data, *x*_ST_ denotes spatial transcriptomic data, and *s*_ST_ denotes spatial coordinates. The superscripts *g* and *s* mark gene expression and spatial information, respectively. The raw data *x*_SC_, *x*_ST_ and s_ST_ in the first step of the pipeline is used to create the spatially annotated data (*x, s*) in step 2 which is then analyzed for the desired statistical inference.

**Figure 2:**
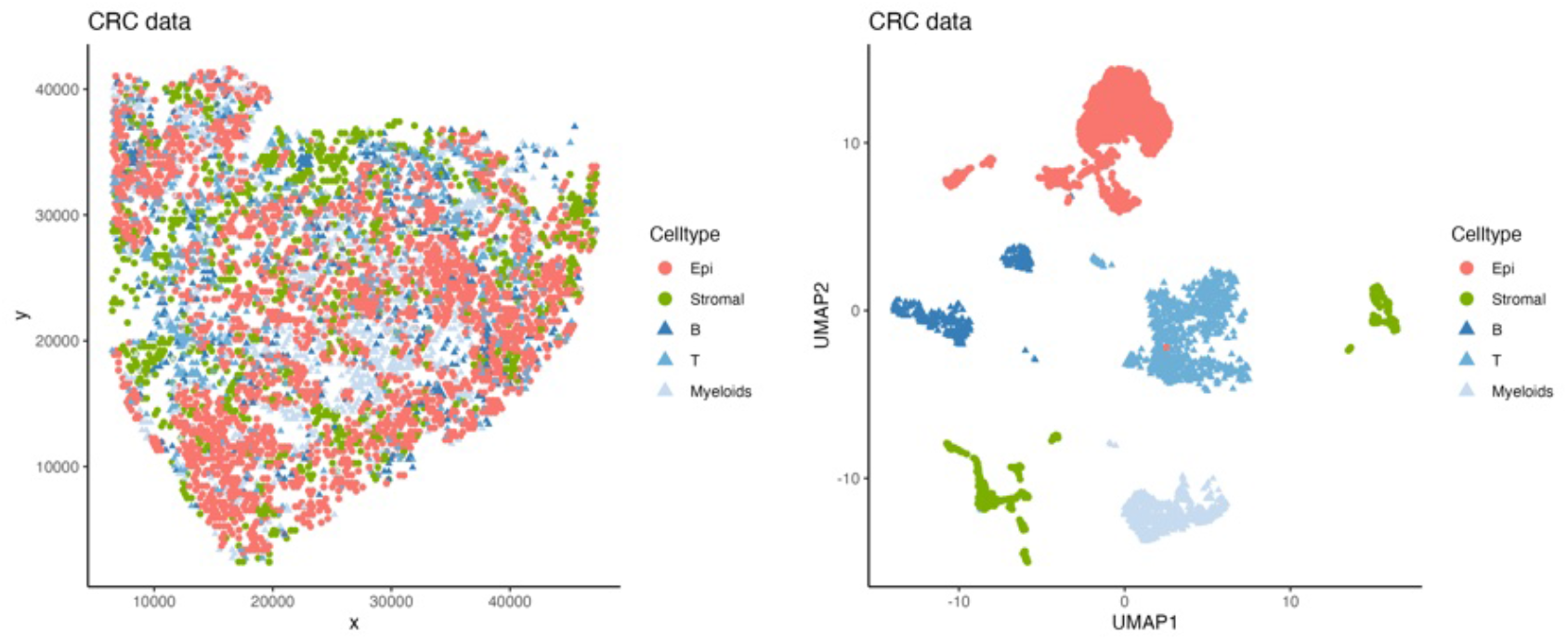
Spatially enriched CRC single cell data after preprocessing, plotted w.r.t. spatial coordinates (left panel) and UMAP scores of gene expression (right panel). B, T and Myeloids refer to immune subpopulations, Epi refers to tumor (epithelial) cells, and Stromal refers to stromal cells.

## 3 A spatially aligned random partition model (SARP)

### 3.1 Statistical model

Through cell mapping and preprocessing of ST and SC data, we obtain tumor, stromal and immune single cell data with imputed spatial locations. We first consider inference for two sets of cells, immune and non-immune cells (the latter including tumor and stromal cells), and defer extension to more than two types to later (Section 3.3). The main analysis goal remains inference on spatially aligned partitions for the two types of experimental units, immune and non-immune cells. For each of the two types, we use clustering to discover subtypes representing homogeneous cell subpopulations. The desired spatial alignment of some of these subtypes will reveal how immune cells interact with subtypes of non-immune cells.

We introduce the construction of the spatially aligned random partition models. Let ***y***_*ji*_ denote the feature vector including gene expression and spatial coordinates for non-immune cells (*j* = 1; *i* = 1, …, *n*_1_) and immune cells (*j* = 2; *i* = 1, …, *n*_2_), respectively. Here *n*_1_ and *n*_2_ denote the numbers of non-immune and immune cells. We assume that the features ***y***_*ji*_ are generated from mixtures ∫ *f*(· | ***θ***) *dG*_*j*_(***θ***) of a kernel *f*(·) with respect to discrete mixing measures *G*_*j*_. We write the mixtures equivalently as hierarchical models with cell-specific latent variables ***θ***_*ji*_, *j* = 1, 2; *i* = 1, …, *n*_*j*_:

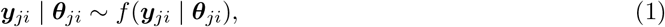

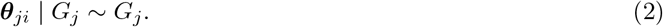

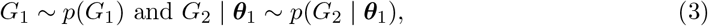

where the conditional prior on *G*_2_ | ***θ***_1_ introduces the desired spatial co-location (details later). The discrete nature of *G*_*j*_ induces ties among the ***θ***_*ji*_ which create a random partition of the experimental units (cells, in this case).

First, consider the special case that ***y***_*ji*_ only contains the spatial coordinates whose distributions should be aligned across subtypes of the two different experimental units. The dependence across types *j* = 1, 2, is introduced by way of the prior (3). We use a construction based on the Pitman-Yor process (PYP), also known as the two-parameter Poisson–Dirichlet process. It is a generalization of the widely used Dirichlet process (DP). For a brief review of the DP and the PYP see, for example, [28]. One of the many defining properties of the DP is the following. A random probability measure is said to follow a DP with concentration parameter *a* and base measure *G*_0_, *G* ~ DP(*a, G*_0_), if for any finite partition *A*_1_,…, *A*_*k*_ of the sample space, the random variables *G*(*A*_1_),…, *G*(*A*_*k*_) follow a Dirichlet distribution with parameters (*a G*_0_(*A*_1_),…, *a G*_0_(*A*_*k*_)). Another defining property of the DP is linked to the related random partition. The random probability measure *G* is almost surely discrete, implying positive probability for ties in a random sample 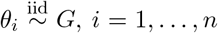, for *G* ~ DP(*a, G*_0_). The DP can be characterized via the random partition that is implied by the configuration of such ties. The law of this random partition is known as the Chinese restaurant process (CRP). Under the CRP the probability for the (*n* + 1)-st observation joining an existing cluster *h* is proportional to *n*_*h*_, where *n*_*h*_ is the number of observations in cluster *h*, and the probability of creating a new cluster is proportional to *a*. The PYP is a generalization of the DP by introducing a discount parameter *σ*. The induced partition structure follows the CRP with an additional discount parameter, with the probability of assigning the (*n* + 1)-st observation to an existing cluster *h* is proportional to *n*_*h*_ − *σ*, and the probability of creating a new cluster is proportional to *a* + *σH*, where *H* is the total number of clusters.

We then specify the prior (3) by assuming *G*_1_ to be generated from a Pitman-Yor process (PYP) with non-atomic base measure 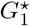. And conditional on parameters *θ*_1*i*_ ~ *G*_1_, *i* = 1,…, *n*_1_, the mixing measure *G*_2_ is assumed to be generated from another PYP, but now conditional on 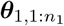:

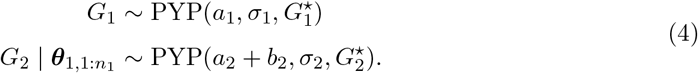

While we consider only spatial coordinates we use 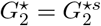 (with a superindex ^*s*^ in anticipation of later model extension beyond spatial features). The construction of 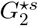 is where we introduce the desired dependence by allowing the atoms of 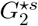 to emerge at locations around ***θ***_1*i*_, *i* = 1, …, *n*_1_, or at new locations. Specifically, the base measure 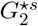 is a mixture of a new base measure 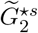 and kernels centered at 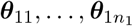, with relative weights *a*_2_ and *b*_2_,

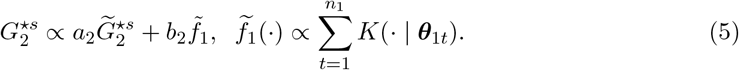

This construction makes *G*_2_ dependent on 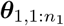, which explicitly models the connection between clusters of two types of cells. Note that the formulation of *G*_2_ is dependent on *n*_1_. This breaks Kolmogorov consistency. That is, in general, the implied model for (*n*_1_ − 1, *n*_2_) cells of the two types is not the marginal of the same for (*n*_1_, *n*_2_) units. However, as we shall briefly discuss later, there is a special case when (1)-(5) could be characterized as a DP mixture for the merged set of all *n*_1_ + *n*_2_ units. In that case Kolmogorov consistency is recovered.

In the application to clustering cells, we set *σ*_1_, *σ*_2_ < 0 to imply a priori a moderate number of clusters for the cell subtypes, favoring larger clusters with reasonable biological interpretations. The proposed model uses *a*_1_, *a*_2_, *b*_2_ and different base measures 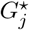 to allow control and flexibility of the dependence between conditions. Additionally, we use the same parametric family for *K*(·) as the sampling distribution *f*(·) in (1). This simplifies computation by allowing to treat the atoms in 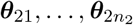 that are generated from *K*(· | ***θ***_1*i*_) as if they were additional observations, and leads to convenient posterior simulation schemes (details in Section 3.4). In general *K*(·) should be chosen to match a scale that would allow biologically plausible cell-cell communication.

Next we extend the model to the full observed feature vectors ***y***_*ji*_ = (***s***_*ji*_, ***x***_*ji*_), *j* = 1, 2, including *D*−dimensional (typically *D* = 2) spatial coordinates ***s***_*ji*_ and the *G*− dimensional gene expression features ***x***_*ji*_. Dependence across the two types is restricted to the spatial subvector by the following construction. Recall that the aim is to represent co-location of two different types of cells. There is no biologic reason why, for example, immune cell gene expression should be correlated with tumor cell gene expression. This is why we restrict the correlation to the spatial subvector. The model construction proceeds as follows. We assume independent kernels on gene expression features and spatial locations, denoted as *f*_*x*_ and *f*_*s*_, respectively,

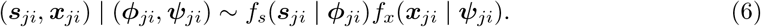

Here the parameter vector ***θ***_*ji*_ = **(*ϕ***_*ji*_, ***ψ***_*ji*_) ∈ 𝒮 × 𝒳_*j*_ with ***ϕ***_*ji*_ indexing the sampling model for the spatial variable, and ***ψ***_*ji*_ indexing the model for gene expression for cell types *j*. We assume independence of **(*ϕ***_*ji*_, ***ψ***_*ji*_) in base measures 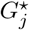,

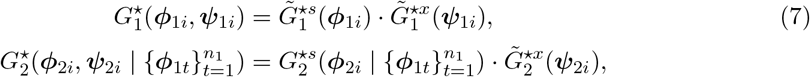

where the prior distribution for the spatial subvector ***ϕ***_2*i*_ in 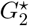 includes the desired spatial dependence structure as in (5). The sampling model (6) replaces (1). Together with (2), (4) and (7) this completes the structure of the proposed spatially aligned random partition models (SARP). See below for specific parametric assumptions.

Specifically, in our application, we assume independent Gaussian distributions for spatial coordinates and gene expressions in the sampling model, such that,

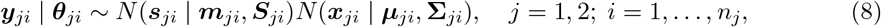

with spatial parameters ***ϕ***_*ji*_ = (***m***_*ji*_, ***S***_*ji*_) and gene expression parameters ***ψ***_*ji*_ = (***µ***_*ji*_, **∑**_*ji*_). We factor ***S***_*ji*_ as ***S***_*ji*_ = **Γ**_*ji*_***R***_*ji*_ **Γ**_*ji*_, with standard deviations **Γ**_*ji*_ = diag(*γ*_*ji*1_,…, *γ*_*jiD*_) and correlation matrix ***R***_*ji*_, to allow for more control over the spatial profiles of the cell sub-populations. The PYP base measures 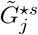 in (7) related to the spatial feature parameters ***ϕ***_*ji*_ = (***m***_*ji*_, **Γ**_*ji*_, ***R***_*ji*_) are given by

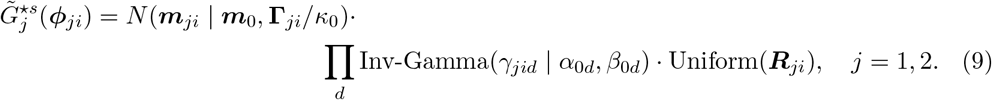

In the case of bivariate spatial coordinates the uniform on ***R***_*ji*_ reduces to a uniform prior on (−1, 1) on the scalar correlation coefficient. For the kernel *K*(· | ***ϕ***_1*t*_) in (5) we assume the same form as 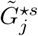 in Equation (9), replacing ***m***_0_ by ***m***_1*t*_. We suggest using informative priors on the variance parameters *γ*_*jid*_ in (9) so that the reported co-localized clusters are within a radius of each other for biologically plausible cell-cell communication and therefore meaningful interpretation of the aligned clusters. For gene expression features, we assume normal inverse-Wishart (NIW) base measures

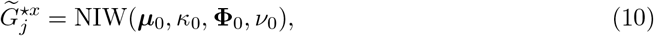

for *j* = 1, 2. The inverse gamma prior on *γ* in (9) ensures that implied clusters remain spatially localized. Without a separation of the prior specification on ***ϕ*** and ***ψ***, cluster formation could be unduly dominated by high-dimensional gene expression features.

In summary, the SARP model introduces an aligned random partition model for non-immune and immune cells, characterizing cell subtypes with homogeneous spatial and gene expression profiles. The alignment is formalized as dependence in the spatial coordinate sub-vector. Figure 3 shows a diagram of the SARP model structure. Note the inherent asymmetry in the model, reflecting the asymmetric, causal nature of the underlying biology with tumor cells recruiting immune cell subpopulations. The involvement of two (or more) sets of experimental units of different nature, with dependence only arising in a subvector of the features, sets the SARP construction apart from similar constructions of dependent BNP priors for dependent mixture models, for example, in [18, 29]. Finally, the very different nature of the two sets of experimental units is also reflected in the choice of kernels. Immune cells have different gene expression profiles from non-immune cells even when they are spatially aligned. There is no notion of the dependence extending to the gene expression part of the data vectors.

**Figure 3:**
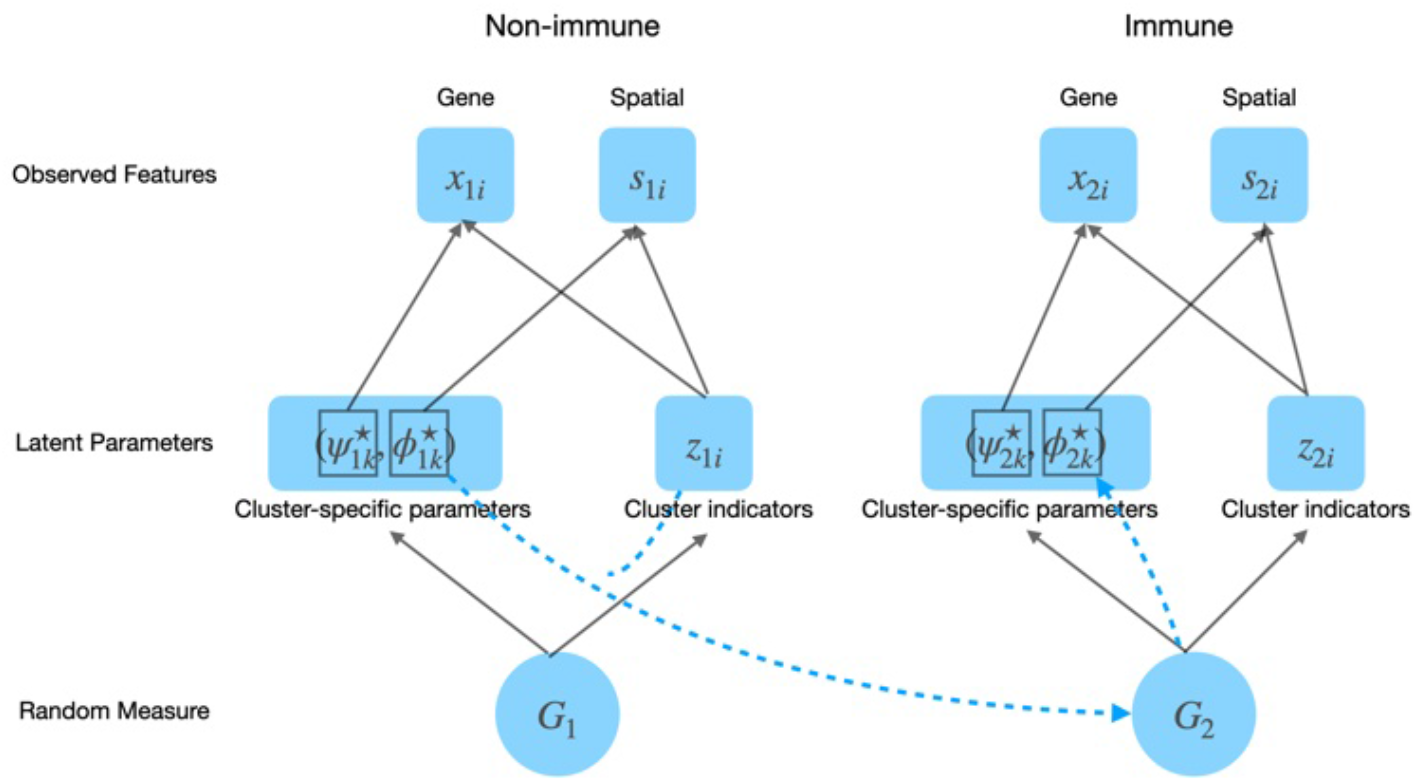
Diagram of SARP model on non-immune and immune cells. The blue dashed lines illustrate the dependency structure for spatial alignment between immune and non-immune cells. The arrows originating from 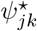 and 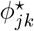 indicate the corresponding observed features.

### 3.2 Properties

We discuss some properties of the SARP model. We first introduce an alternative approach for inference that clusters the pooled data including both immune and non-immune cells. The model on the joint data is shown to be a special case of our proposed model under specific hyperparameter settings. This illustrates the roles of hyper-parameters and the proposed structure in the SARP model. Next we give the posterior contraction rate of the mixing measures in the SARP model. Lastly, we characterize the probability function of the random partitions induced by the SARP model. All discussed results are based on well known results applied to the specific model only. Brief proofs are provided in Appendix A of the supplementary material.

We first discuss the joint distribution of the spatial features ***s***_*ji*_ across both types of cells under the SARP model, focusing on the spatial subvector for the moment. Choosing *K*(· | ***ϕ***) = *δ*_***ϕ***_(·) in (5) the distribution 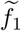 reduces to the empirical distribution of 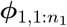. Under this assumption and the special case of the DP prior with discount parameters *σ*_1_ = *σ*_2_ = 0, and if *a*_1_, *a*_2_ and *b*_2_ are chosen accordingly, then (***ϕ***_*ji*_; *j* = 1, 2 and *i* = 1, …, *n*_*j*_) arise as an i.i.d. sample from a single mixing measure *G* with a DP prior. The result clarifies the model structure by recognizing the SARP model as breaking pooled data clusters into co-localized non-immune and immune cell clusters, with *K*(· | ***ϕ***) ≠ *δ*_***ϕ***_ allowing some dis-location of the latter. If *σ*_1_ = *σ*_2_ = 0, *a*_2_ = *a*_1_, *b*_2_ = *n*_1_, *K*(· | ***ϕ***) = *δ*_***ϕ***_(·), and 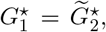, the model in equations (2), (4) and (5) is equivalent to

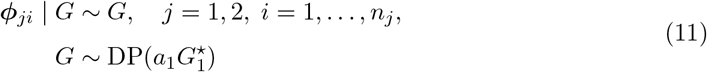

The SARP model introduces latent ***θ***_*j,i*_ generated from discrete random measures with PYP priors as specified in (4). Instead of using the PYP prior, one could consider any suitable Bayesian nonparametric prior, incorporating the same dependence structure on the base measures. For example, finite mixtures have proven useful in many applications. In fact, a PYP prior *G*_*j*_ ~ PYP(*M*_*j*_, *σ*_*j*_, *G*^✶^) with negative discounting parameters *σ*_*j*_ < 0 and total mass parameters *M*_*j*_ = −*Kσ*_*j*_, *K* ∈ {2, 3,…} is equivalent to a finite mixture model with weights from a symmetric Dirichlet distribution Dir(−*σ*_*j*_,…, −*σ*_*j*_) and atoms from the base measure *G*^✶^ [7, 30]. More generally, with a prior on the hyper parameter *K* the model becomes a mixture of finite mixtures (MFM). In a MFM, the latent parameters ***θ***_*ji*_ are modelled as ***θ***_*ji*_ | *K*, 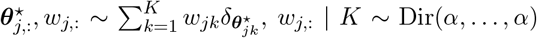, and *K* ~ *p*_*K*_(*K*) where where *p*_*K*_ is a prior on the number of components. Using a MFM we can characterize posterior asymptotics. Under the MFM prior, the posterior distribution 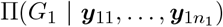 and the conditional posterior distribution 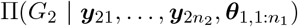 converge with increasing *n*_*j*_ to the true mixing measure 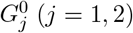, with posterior contraction rates as follows.

Under model (8) and MFM prior, for *j* = 1, 2, we have

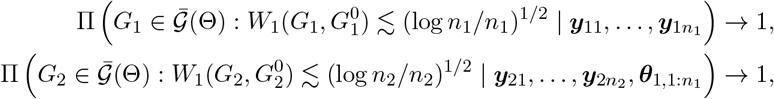

in probability under 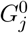. Here, 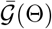 denotes the space of all discrete measures, and *W*_1_ denotes *L*_1_ Wasserstein distance. The relation ≲ denotes inequality up to a constant multiple where the value of the constant is independent of *n*_*j*_. The results are for the random partition for each cell type, as *n*_*j*_ → ∞, keeping the sample size of the other cell type fixed. The result remains valid under a PYP prior with *σ*_*j*_ < 0 and *M*_*j*_ = −*Kσ*_*j*_, *K* ∈ {2, 3,…}, as we use it in our implementation, assuming that *K* equals to the true number of clusters. This follows from the result in Equation (4) of [31]. Extension to general PYP priors with positive discount parameters is beyond the scope of this paper, as the cited results in [32] are based on finite mixtures and DP mixtures.

The proposed SARP model introduces random partitions of cells, with clusters of cells being interpreted as cell subtypes. Let *M*_1_ = *a*_1_ and *M*_2_ = *a*_2_ + *b*_2_ denote the total mass parameter of the two PYPs in Equation (4). Using hyperparameters (*σ*_*j*_ < 0 and *M*_*j*_ ∈ {−2*σ*, −3*σ*, …}) or (*σ*_*j*_ ∈ [0, 1) and *M*_*j*_ > −*σ*_*j*_), the implied random partitions can be described as follows. Let 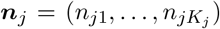 denote the cardinality of *K*_*j*_ clusters of the *N*_*j*_ = ∑ _*ℓ*_ *n*_*jσ*_ samples from group *j, j* = 1, 2. The marginal prior for the random partition ***n***_*j*_ under the PYP priors in (4) is then 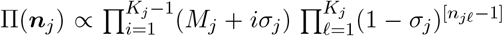 [7]. As the random partition implied by i.i.d. sampling from *G*_*j*_ the model ∏(***n***_*j*_) preserves Kolmogorov consistency (across sample sizes). However, the joint model *p*(***n***_1_, ***n***_2_) implied by 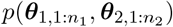 loses this coherence across sample size, except for the special case exhibited before, in (11). In summary, the SARP model introduces structure to allow the desired inference on spatial alignment in the application, at the price of breaking the coherence property.

### 3.3 Extension to multiple cell types

In the previous discussions, we split the dataset into immune versus non-immune cells, and investigated the subtypes of immune cells that are recruited by certain non-immune subtypes. A natural extension of the model is then to more than two types of cells. In the application to the CRC data, we are interested in three types of cells, including tumor, stromal, and immune cells, and investigate how subpopulations of the latter two cell types are co-located with tumor cell sub-types. We therefore generalize the SARP model to spatially aligned clustering of multiple (≥ 2) types of experimental units where one of the types of units is considered as reference for the spatial co-localization.

Let *j* = 1, …, *J* index *J* types of experimental units. Without loss of generality, assume *j* = 1 is the reference type of units, such that some clusters of type *j* (*j* > 1) units should locate around clusters of type 1 units. In our case, *j* = 1, 2, 3, denotes tumor, stromal, and immune cells, respectively. We follow the same construction as in Section 3.1. The sampling model and prior on *θ* remain as in (1) and (2), and we specify a joint prior on 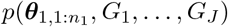 by assuming independence of *G*_*j*_ (*j* > 1) given 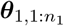, i.e., 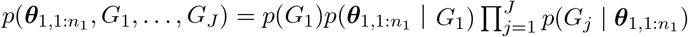, with

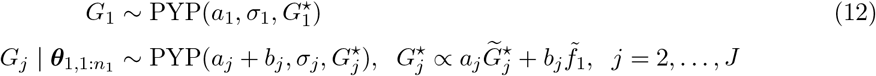

With 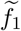 as in (5), and the base measures 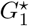 and 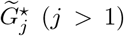 are specified like 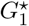 and 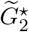 before, and hyperparameters remain unchanged. Under *J* = 2 the model reduces to the earlier construction.

### 3.4 Posterior Inference

Posterior inference is implemented as MCMC simulation. We introduce latent cluster membership indicators *z*_*ji*_ with 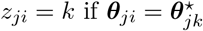, where 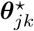, *k* = 1, …, *K*_*j*_ are the unique elements of 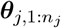. We can rewrite (1) as

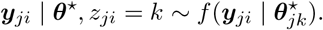

Under the PYP the implied prior for the indicators *z*_*ji*_ can be characterized as *p*(*z*_*ji*_ = *k*) = *w*_*jk*_ with *w*_*jk*_ generated by a stick breaking construction, *w*_*j*1_ = *V*_*j*1_, 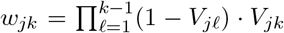, and *V*_*jk*_ ~ Beta(1 − *σ*_*j*_, *M*_*j*_ + k*σ*_*j*_), where *M*_1_ = *a*_1_, *M*_*j*_ = *a*_*j*_ + *b*_*j*_, *j* = 2, …, *J*. The prior models for the atoms 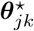 are

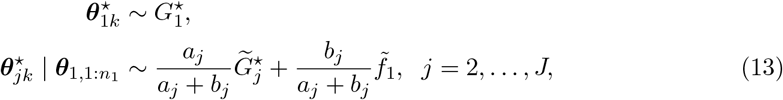

where 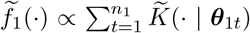. Here we rewrite the model with 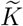 defined for ***θ*** = (***ϕ, ψ***) for a convenient notation in the MCMC algorithm. Defining 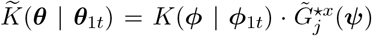 the model remains unchanged. We introduce additional indicators *c*_*jk*_ ∈ {0, 1,…, *n*_1_}, *k* = 1,…, *K*_*j*_ to replace (13) by

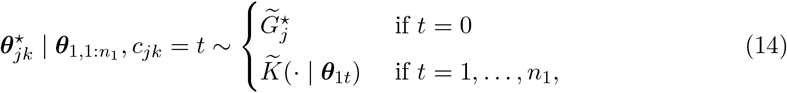

with

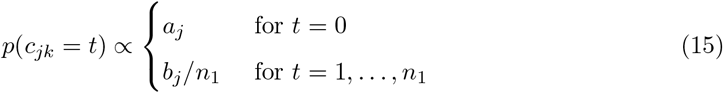

For a better mixing Markov chain we use a blocked Gibbs sampler based on a finite truncation of the PYP random probability measure, truncating at *K* = *K*_*max*_. Note that the posterior inference algorithm in this section is applicable for any general choice of hyper parameter in PYPs. The truncation *K* = *K*_*max*_ is only needed when *σ*_*j*_ > 0, whereas for *σ*_*j*_ < 0 the model becomes a finite mixture as discussed in Section 3.2 and the number of components *K* is given by −*M*_*j*_/*σ*_*j*_ where *M*_*j*_ is the total mass parameter in PYP. Using the indicators *c*_*jk*_ and *z*_*ij*_ the full conditional posterior probabilities can be derived in closed form. This motivates the following Gibbs sampling transition probabilities to define MCMC posterior simulation:

1. Update ***θ***^⋆^ by generating from the complete conditional posteriors. Let *L*_*jk*_ = {*ℓ* : *c*_*jℓ*_ = *i, z*_1*i*_ = *k*}. Then

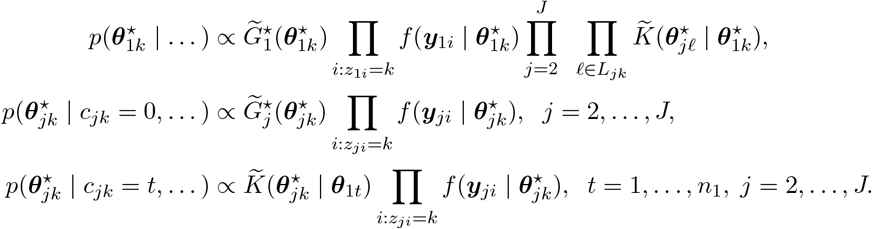

2. Update *z*_1*i*_ for *i* = 1, …, *n*_1_, using

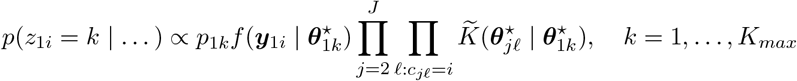

3. Update *z*_*ji*_ for *i* = 1, …, *n*_*j*_, *j* = 2, …, *J*, using

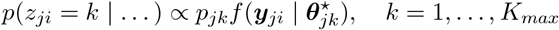

4. Update V_*j,k*_ for *j* = 1, …, *J, k* = 1,…,*K*_*max*_ − 1. Let 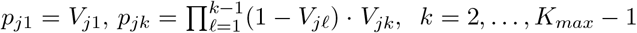. Sample

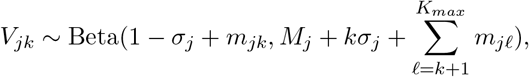

where *m*_*jk*_ is the cardinality of {*i* : *z*_*ji*_ = *k*}.

5. Update *c*_*jk*_ for all currently imputed clusters, *j* = 2, …, *J*,

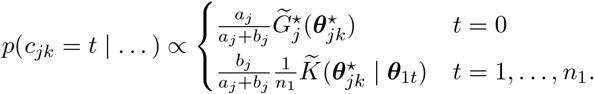

Based on the conjugate prior assumptions all complete conditional posterior distributions are available in closed form and can be easily sampled from. Details are given in Appendix B.

## 4 Simulation Study

We carry out simulation studies to assess inference under the proposed model. Experiments include analyses with simulated data, and with semi-synthetic data. Simulated data allows to verify the ability to discover spatial alignment under realistic assumptions on the true partition. Using simplified bivariate gene expression features we can assess inference by visualization. Simulation with semi-synthetic data based on real single-cell data implments evaluation of inference on co-location under actually observed higher dimensional marginal distributions. In all simulation studies we focus on two types of cells, such as immune versus non-immune cells.

### Hyper-parameters

Throughout the simulation studies, we implement the SARP model with parametric families as specified in Section 3.1. For the hyper-parameters in PYP priors, we use *a*_1_ = 2, *a*_*j*_ = 1, *b*_*j*_ = 1, *j* = 2, …, *J*, and discount parameters *σ*_*j*_ = −2/*K*_*max*_ where *K*_*max*_ = 10. The hyper-parameters in the base measures are set as follows: for the parameters on spatial features in Equation (9), ***m***_0_ is the sample mean, *κ*_0_ = 1, *α*_0*d*_ = 10^5^, *β*_0*d*_ = (sample variance)/100*α*_0*d*_. For the parameters on gene expression in Equation (10), ***µ***_0_ is the sample mean, **Φ**_0_/100 is the sample covariance matrix, *κ*_0_ = 1, *ν*_0_ = 2 + (dimension of gene expressions).

### Comparison

We compare results under SARP with inference under a Gaussian mixture model (GMM) as a benchmark. The number of clusters for the GMM is determined based on an ELBOW plot. In the semi-simulated study we also compared with SpaRTaCo [4]. SpaRTaCo is added in the comparison since in contrast to the GMM it does perform co-clustering based on both spatial and gene-expression features, making it conceptually closer to the proposed model (although co-location is not explicitly considered). Since GMM and SpaRTaCo do not include explicit parameters to indicate co-localization we have to add a suitable posterior summary. Given the discovered clusters under the GMM, for the purpose of summarizing simulation results we define the probability of an immune cluster *k*_2_ being co-localized with a non-immune cluster *k*_1_ (or spatially independent) as proportional to the likelihood of the immune cells in cluster *k*_2_ under the kernel with cluster-specific parameters of non-immune cluster *k*_1_ (or the distribution used in 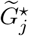). Such cluster-specific co-localization probabilities naturally imply cell-cell co-localization probabilities to be used in the following discussion of reports.

Reporting and summarizing inference on the random partition (*z*_*ji*_) and on the co-localization of clusters (*c*_2*j*_) requires some more clarifications.

### Random partition

We report a posterior summary of the random partition by minimizing the variation of information (VI) loss as proposed in Dahl et al. [33], and implemented in the *salso* R package. The reported point estimate 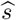 of the random partition characterizes subpopulations of immune and non-immune cells.

### Co-location

We define co-localization summaries at the cluster and at the cell-level under the simulation truth and under the inference, using notation with a [*K*_1_ × *K*_2_] matrix ***P*** for cluster-level summaries, and a [*n*_1_ × *n*_2_] matrix ***Q*** for cell-level summaries (and [*K*_2_ × *n*_1_] ***PQ*** for a mixed cluster-cell summary – see below), and a superindex ^*o*^ to mark the simulation truth. For each immune cell cluster (*j* = 2, *k*) and non-immune *cell t*, the posterior probabilities 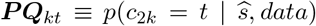 summarize inference on co-localization, including characterization of uncertainties. Note that following the structure of the inference model ***PQ***_*kt*_ is cell-specific for non-immune cell *t*. Alternatively, a cluster-specific posterior probability of co-localization for immune and non-immune clusters *k* and *ℓ*, respectively, can be reported as 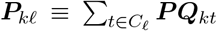, with *C*_*ℓ*_ = {*t* : *z*_1*t*_ = ℓ}. A limitation of these summaries is the conditioning on cluster labels. Instead, we report a summary that avoids label-switching by focusing on cell-specific co-localization. Let ***Q*** denote an (*n*_2_ × *n*_1_) cell by cell co-localization matrix with 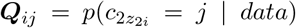, the probability that immune cell *i* appears in a cluster that is linked with non-immune cell *j*. The probability of immune cell *i* being co-localized with any of the non-immune clusters is represented by the row sums of matrix ***Q***, i.e. *q*_*i*_ = ∑_*j*_ ***Q***_*ij*_. We will define a (*K*_1_ × *K*_2_) true binary co-localization status matrix ***P*** ^*o*^ between immune and the non-immune clusters under the ground truth (naturally ***P*** ^*o*^ is a binary matrix), which induces a (*n*_2_ × *n*_1_) true cell-cell co-localization matrix ***Q***^*o*^ (details when we introduce the simulation scenarios). Also 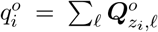 is an indicator that cell *i* is co-localized with a non-immune cluster under the ground truth. We report the accuracy of estimating 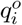 as 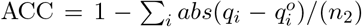. For a practically more relevant summary of cell-cell co-localization, we report true negative rate (TNR, specificity) and true positive rate (TPR, sensitivity) as a function of a threshold *λ* on ***Q***_*ij*_ as follows. Let 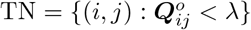 and *N*_0_ = |TN|, and similarly 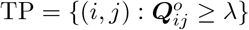 and *N*_1_ = |TP|. Let then TPR = ∑_TP_*I*(***Q***_*ij*_ ≥ *λ*) *N*_1_, and TNR = ∑ *I*(***Q***_*ij*_ < *λ*) *N*_0_. Accordingly, the correct classification rate is TCR = {∑ TP *I*(***Q***_*ij*_ ≥ *λ*)+ ∑TN *I*(***Q***_*ij*_ < *λ*)} (*N*_1_ + *N*_0_). In the simulation results we chose λ to achieve a set TNR, and then compare the corresponding TPR and TCR.

### 4.1 Simulated data

We simulate data assuming bivariate gene expression features ***x***_*ji*_ and spatial coordinates ***s***_*ji*_, mainly for the ease of visualization. The first row of Figure 4 shows the simulated cells plotting spatial coordinates and gene expression, respectively. Considering three true non-immune cell clusters, 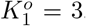, and four immune cell clusters, 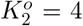, we first specify a ground truth for the co-localization between the immune and non-immune clusters as indicated in the first row of Table 1. The first three immune clusters are co-localized with non-immune cluster 2,2,1, respectively, and the fourth cluster is spatially independent. This defines the true cluster-cluster indicators ***P*** ^*o*^. From ***P*** ^*o*^, we induce a cell-cell co-localization truth ***Q***^*o*^ by spreading any entry 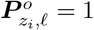 for immune cell *i* uniformly across all non-immune cells *j* in cluster *ℓ* (to be used for the reported comparisons). We then specify a Gaussian mixture model with sampling distributions as assumed in (8) with location parameters of the co-located clusters being close to each other spatially, and then generate the bivariate gene expression features as shown in Figure 4ab.

**Table 1:**
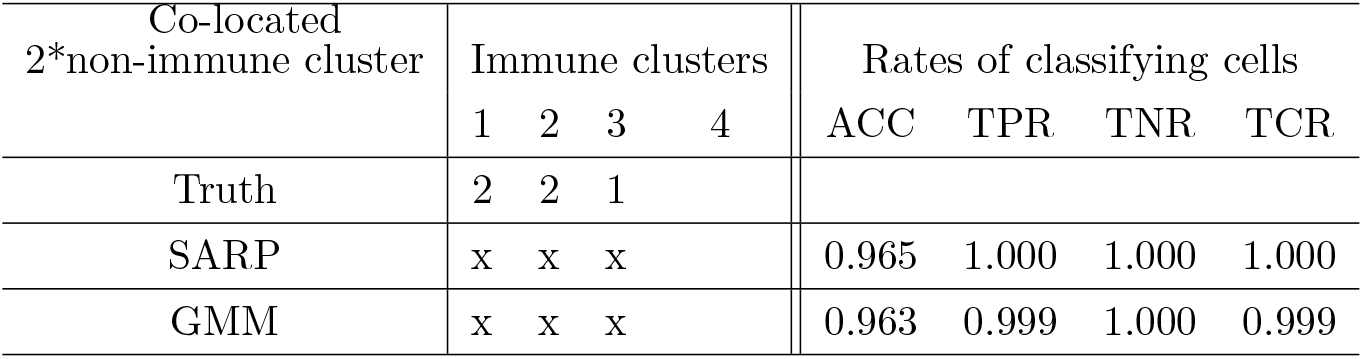
Co-localization between clusters in simulated bivariate data. The symbol “x” denotes the immune cluster is co-localized with one of the non-immune clusters. Blank entries indicate absence of spatial correlation with any non-immune cluster. The specific indices of the linked clusters can not be compared, due to label switching. Instead the last four columns report the accuracy of estimating the cell-by-cell co-location matrix *Q*. See the text for a definition of the rates.

| Co-located<br>2*non-immune cluster | Immune clusters |  |  |  | Rates of classifying cells |  |  |  |
| --- | --- | --- | --- | --- | --- | --- | --- | --- |
|  | 1 | 2 | 3 | 4 | ACC | TPR | TNR | TCR |
| Truth | 2 | 2 | 1 |  |  |  |  |  |
| SARP | x | x | x |  | 0.965 | 1.000 | 1.000 | 1.000 |
| GMM | x | x | x |  | 0.963 | 0.999 | 1.000 | 0.999 |

**Figure 4.**
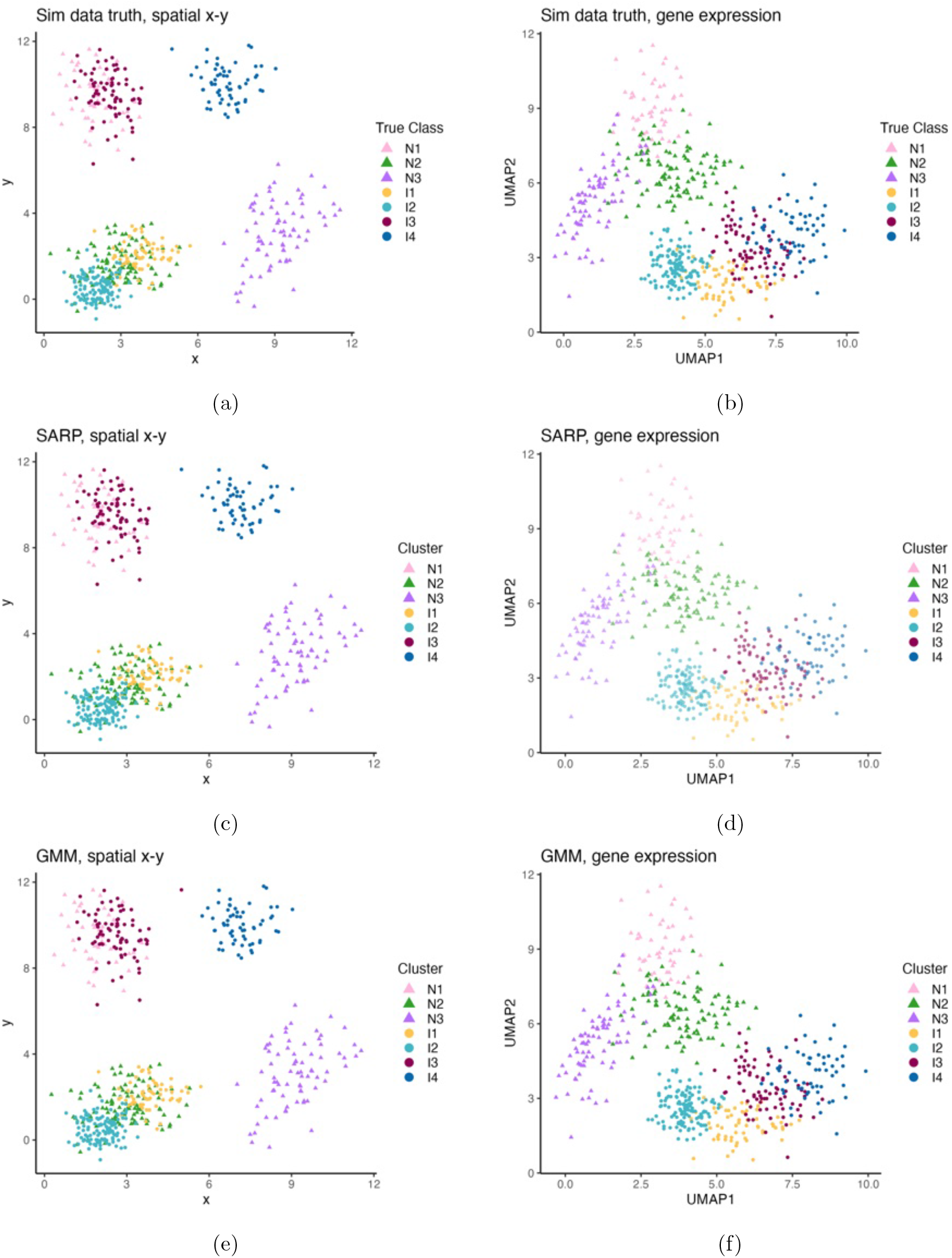
Simulation truth (panels a,b) and estimated partitions under the SARP (c,d) and GMM model (e,f) on 2D data. In the left column the cells are plotted w.r.t. spatial coordinates and in the right column w.r.t. gene expressions.

Implementing inference under the SARP and GMM models for this simulated data, we find an estimate for the random partition. The implied clusters are shown in Figures 4c-f. Table 1 summarizes estimated co-localization of the identified cluster and the accuracy of co-localization under SARP and GMM, respectively. In summary, under this easy scenario with spatially well-separated clusters under the simulation truth, both methods report accurate inference.

### 4.2 Semi-simulated data based on CRC

We generate semi-synthetic data based on the spatially enriched single cell data shown in Section 2 as a reference. In short, we extract the following quantities from the actual data: all cell gene expressions, non-immune cell spatial locations, immune cell cluster assignments by K-means and residuals related to spatial locations; and simulate the remaining ones. The simulation proceeds along the following line: we start with K-means estimates of cluster arrangements for immune and non-immune cells. To create the simulation truth we replace the immune cell cluster centroids 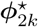 by simulating cluster specific parameters 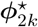 under (14) and (15): we (i) use the estimated 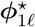; (ii) simulate co-localization indicators 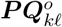; and (iii) generate 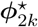 under (15). In more detail, using PCA we get ***x***_*ji*_ ∈ ℜ^*p*^ for *p* = 20 gene expression features, and use the following procedure to generate the semi-simulated data. Using the non-immune cells from the reference dataset we implement K-means to generate a ground truth for non-immune cell clusters, including corresponding cluster centers ***ϕ***_1,*i*_. The K-means algorithm is then also applied to the immune cells. We only save the number of clusters, *K*_2_, cluster memberships *z*_2,*i*_, and cell-specific residuals ***ϵ***_2,*i*_ ≡ ***s***_2,*i*_ − ***ϕ***_2,*i*_, but generate (new) simulated immune cell cluster centers 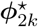 by first randomly generating true co-localization status 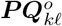, and then, given 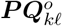 we generate new immune cluster centers 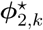 using (14) and (15). Note that the simulation truth on 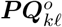 implies a simulation truth 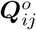 for cell-cell co-localization. Finally, spatial coordinates for immune cells in the semi-synthetic data set are then generated as ***s***_2,*i*_ = ***ϕ***_2,*k*_ + ***ϵ***_2,*i*_ using the earlier saved residuals and cluster centers.

We evaluate the accuracy of identifying co-localization comparing SARP, GMM, and SpaRTaCo. Figure C1 and C2 in the Appendix shows the clustering result and Table 2 summarizes the estimated co-localization between the clusters. Under this semi-simulated scenario that closely mimics the actual data SARP still identifies cell subpopulations and the spatial alignment, even with the relatively high dimensional gene expression and less clearly separated bivariate spatial clusters.

**Table 2:**
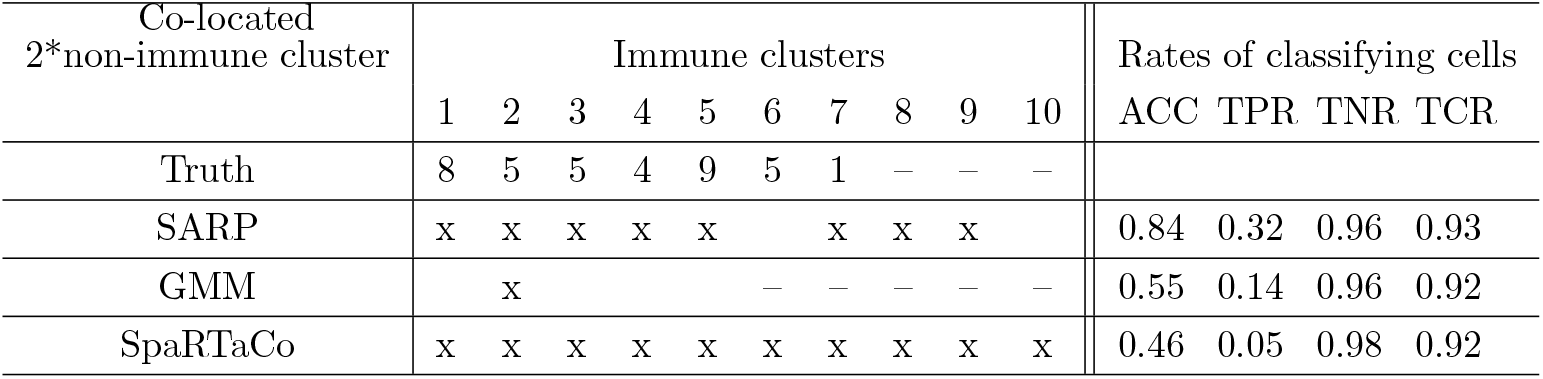
Co-localization across clusters and accuracy in simulated semi-synthetic data. Same as Table 1, for the semi-simulated data.

## 5 CRC data analysis

We fit the proposed SARP model with the CRC data introduced in Section 2, aiming to discover immune and non-immune cell subpopulations, and to understand their spatial alignment. The CRC data (after preprocessing) is shown in Figure 2. In this application the experimental units (indexed by *i*) are the single cells, and their gene expression and inferred spatial locations are the data. For gene expressions ***x***_*ji*_ we use the top 20 PCA dimensions. For the SARP model, we use *K*_*max*_ = 15, and the setting of all other hyper-parameters as described in the simulation studies.

Posterior inference on the partitions reports the following homogeneous cell sub-populations. A point estimate of the partition, i.e., cluster assignment, is obtained as described in the simulation studies. Clusters with closest larger clusters. Note that *K*_*max*_ in the prior specification only sets an upper bound for the number of homogeneous subpopulations that could be meaningfully distinguished.

### Clustering into homogeneous cell sub-populations

The allocation of cells to these clusters can be seen in Figure 5a. We identify 15 non-immune cell clusters and 15 immune cell clusters. We annotate the identified clusters by considering the cell type annotation (using known marker gene as implemented in the annotation under Seurat clustering [34]) of all cells within each cluster, and annotate the identified clusters with the main cell types found in each cluster (clusters are found to be almost pure subsets of the annotated cell types, making the cluster annotation biologically meaningful). The annotated clusters include 11 Epi, 4 Stromal, 6 Myeloid, 7 T-, and 2 B-cell subtypes. The spatial alignment is shown in Figure 5b and we include more discussions on the interpretation with the extended model below. The nature of the reported clusters as almost pure subsets of annotated cell types also validates the choice of *K*_max_ = 15. Larger size partitions would risk to lose the same clean interpretation of clusters as homogeneous cell subpopulations.

**Figure 5.**
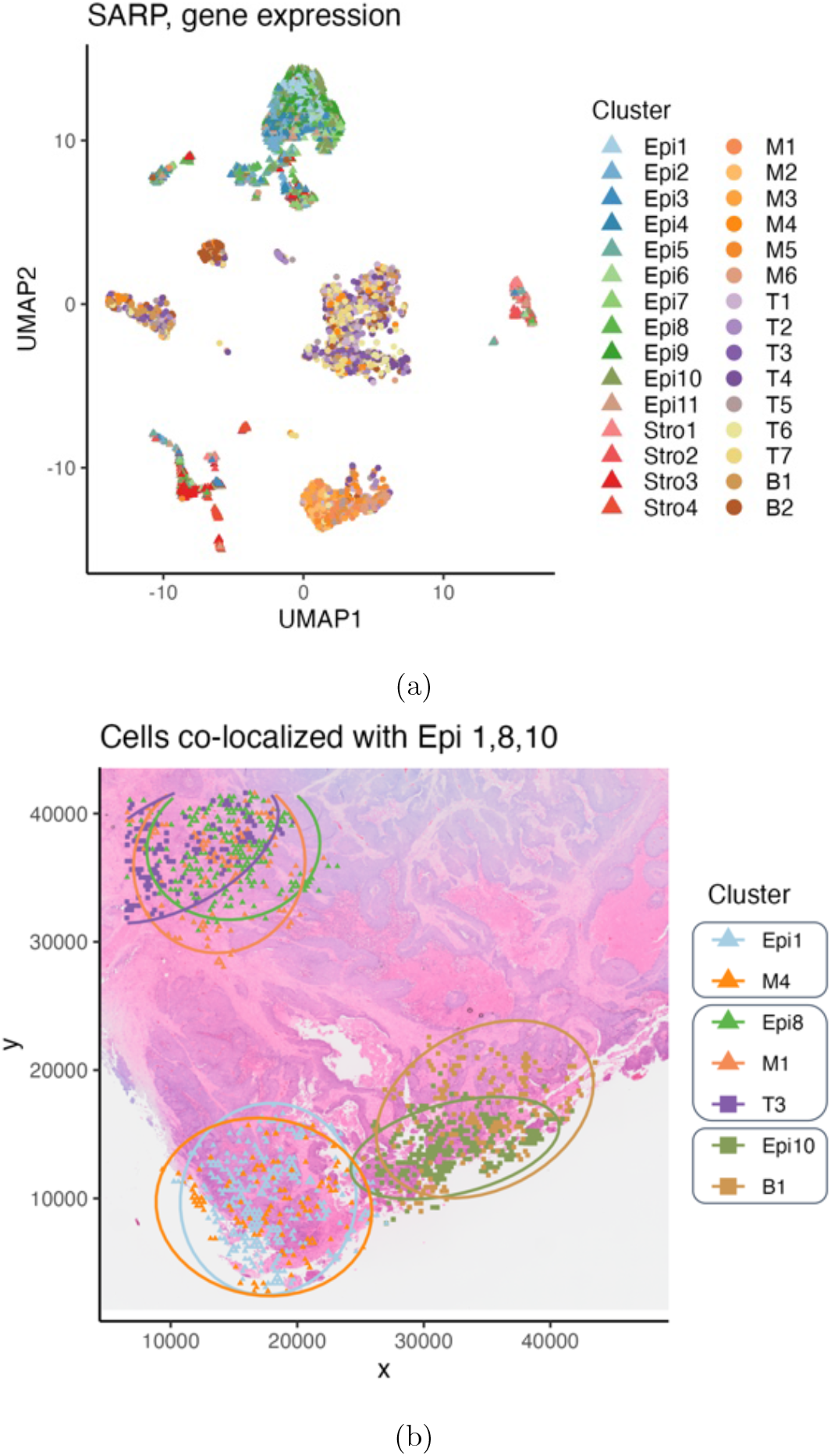
Inference results for the CRC data under the SARP model with two cell types. (a) Estimated clusters plotted w.r.t. UMAP scores of gene expression, colored by estimated clusters. The clusters are indexed by the annotated celltypes. (b) Estimated spatial alignment across the discovered clusters under the 2-type SARP model. The figure show immune cell clusters co-localized with Epi1, Epi8, and Epi10, respectively. All cells are plotted with respect to spatial coordinates. The contours are 95% probability contours of the inferred distribution of the subpopulations on spatial coordinates. For reference, the background shows the histology image. See Figure C1 in appendix C for spatial alignment of the remaining tumor cell clusters.

### Extension to three types

Next we extend the scope of the analysis by considering three types of cells, distinguishing tumor (*j* = 1), immune (*j* = 2), and stromal (*j* = 3) cells. We implement the extended version of the model with *J* = 3 types to investigate co-localization of immune and stromal cell sub-types with tumor subtypes. The setting of the hyperparameters and posterior summaries remains unchanged as in the simulation studies. Inferred cell clusters are shown in Figure 6a. Under the extension to *J* = 3 types we identify a finer resolution of cell sub-populations. Inference under the extended model with *J* = 3 allows more detailed conclusions on how different subtypes of cells interact with tumor cells. The estimated clusters are shown in Figure 6a, and the clusters are annotated using the same approach described in the previous paragraph. We identify 15 epithelial clusters, 14 stromal clusters and 15 immune cell clusters.

**Figure 6.**
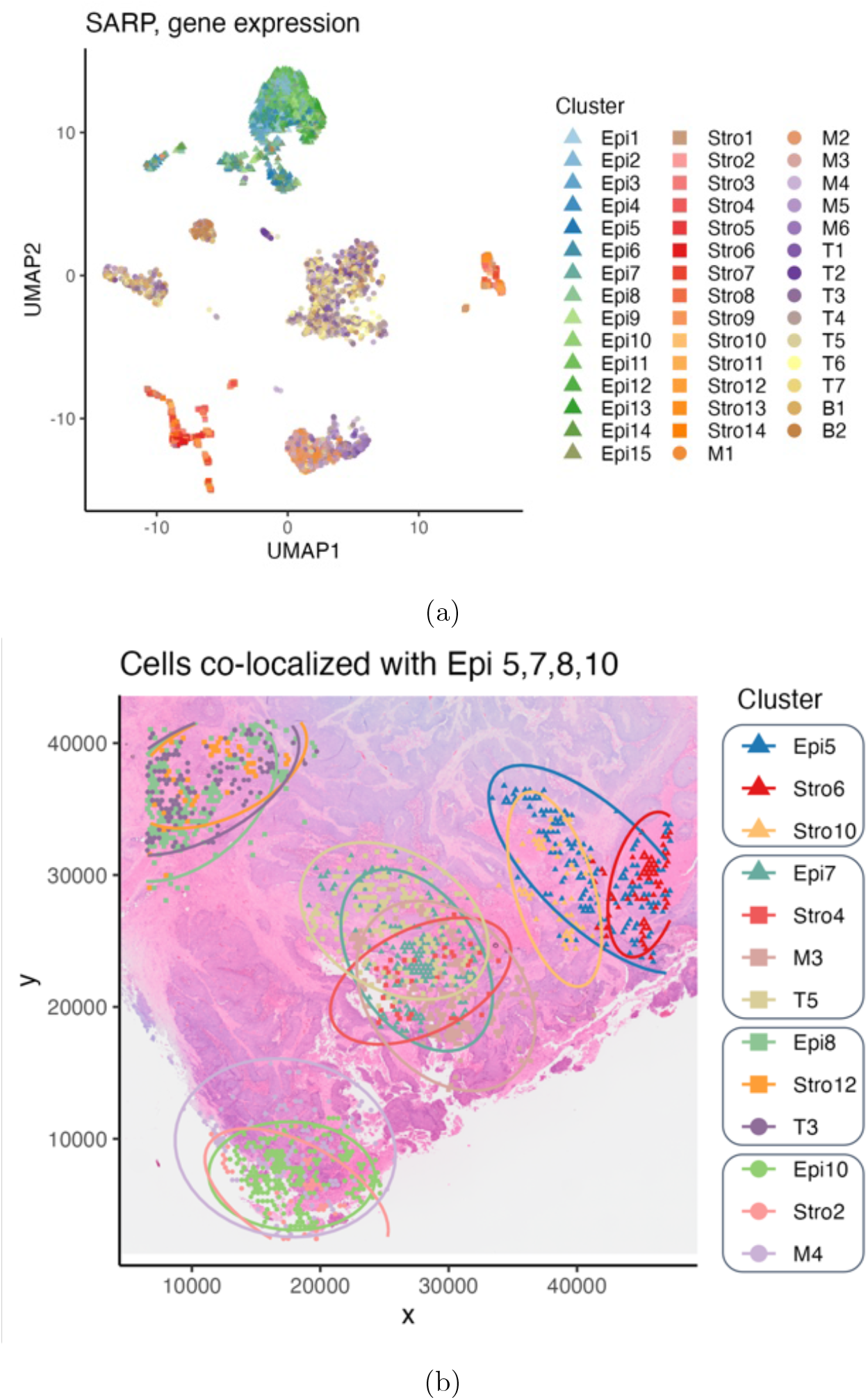
Inference results for the CRC data under the extended SARP model. Same as Figure 5 for the extended SARP model.

### Spatial alignment

Given the cell subtypes, the estimated spatial alignment is shown in Figure 6b. The figure shows the subtypes of immune and stromal cells recruited by different tumor subtypes, including cells that are co-located with epithelial cell subtypes Epi5, 7, 8, 10, respectively. Clusters are plotted w.r.t. the spatial coordinates over the histology image. For example, on the bottom of the figure, we show one myeloid and one stromal cell subtype, are spatially aligned with the epithelial cell subtype 10.

### Uncertainty quantification

Besides point estimates of inferred spatial alignment, uncertainty in the reported co-localization can be reported as follows. For example, immune cluster T4 has a probability of 96% of being spatially aligned with Epi13 (Figure D2 and D3 in the Appendix) and 4% of being not spatially aligned. That is, although sub-population T4 is (*a posteriori*) more likely spatially co-localized with Epi13, there is a remaining 4% probability of being spatially independent. Figure D3 in the Appendix plots the posterior probability of immune cells being spatially independent with tumor cells, and the posterior distribution of the number of clusters, to further characterize posterior uncertainty.

For a more in-depth biological interpretation of the observed co-localization patterns, we performed differential expression (DE) analysis among the spatially defined epithelial subsets and identified distinct markers genes per group (Figure 7). Epithelial cells were categorized based on their spatial enrichment for specific components in the tumor micro environment (TME): Epi-Inner (surrounded predominantly by other epithelial cells), Epi-Mixed-I (in proximity to mixed immune populations), Epi-Myeloid, Epi-T, Epi-B, and Epi-Stromal. We focused on confident associations supported by robust marker gene expression and literature validation. For example, Epi-Stromal cells expressed high levels of *CXCL1/2/3*, a chemokine axis previously shown to mediate epithelial-stromal crosstalk and promote a pro-tumorigenic microenvironment [35]. Epi-Myeloid cells were enriched for *AGR2* gene expression, which encodes a protein implicated in macrophage-epithelial communication that promotes an immunosuppressive niche [36]. Meanwhile, Epi-T cells upregulated *CEACAM1* and *CEACAM6*, two cell adhesion molecules known to inhibit antitumor T cell activity and contribute to immune evasion [37]. These results recapitulate well-established epithelial-stromal/immune interactions, supporting the biological relevance of our spatial classification. Furthermore, our framework enables deeper exploration of epithelial phenotypic diversity shaped by local microenvironments, offering new opportunities to define ecosystem-specific vulnerabilities that have the potential to inform therapeutic targeting.

**Figure 7.**
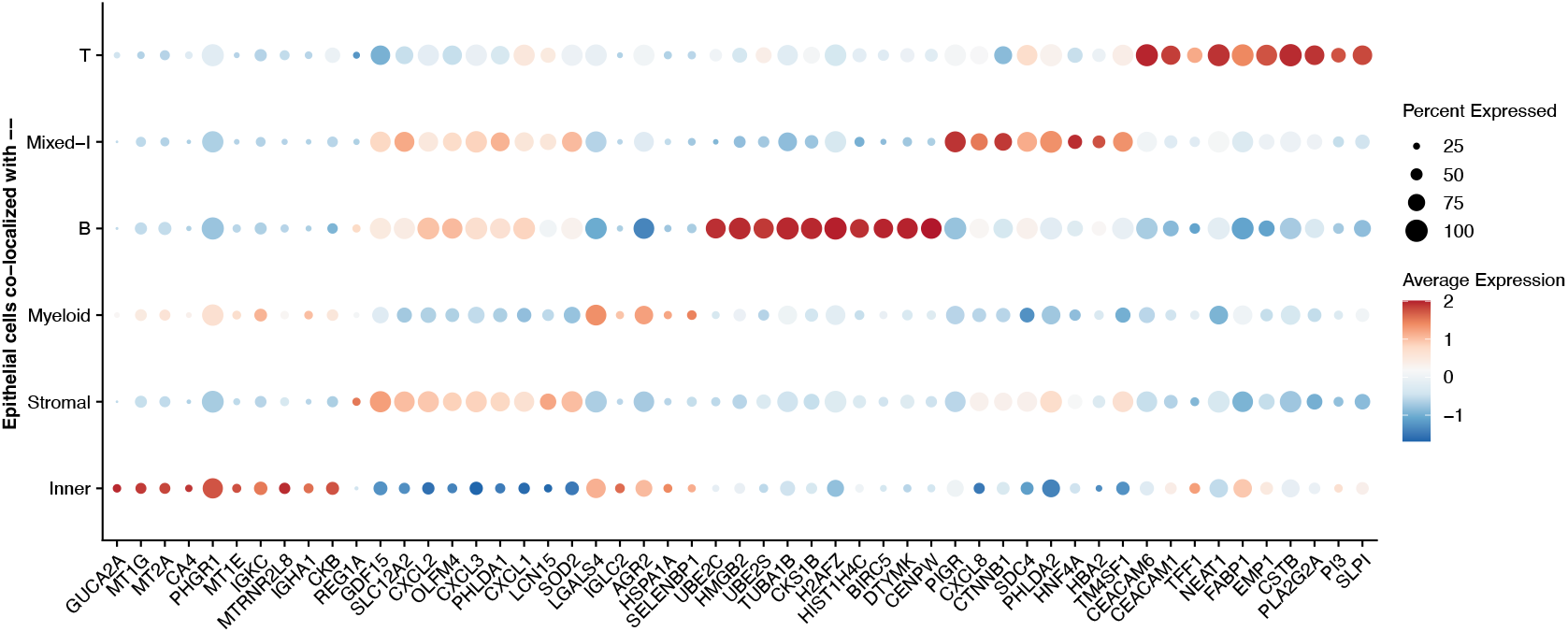
Differential gene expression analysis for epithelial cells by different inferred micro-environment. Epithelial cells are classified into subtypes based on their co-localization with surrounding cells: those recruiting immune cells, including “Myeloid”, “T”, “B”, and “Mixed-I” (mixed immune types); those surrounded by “Stromal” cells; and “Inner” subtypes located within epithelial clusters.

## 6 Discussion

We introduced a spatially aligned random partition model (SARP). SARP addresses a gap in existing methods by introducing inference on regressing clusters within a partition of one type of experimental units to clusters formed by a random partition of another type of experimental unit. Inference under the SARP model identifies clusters on multiple sets of experimental units allowing alignment on a subvector of spatial features, but independence with respect to other features. While we focus on spatial alignment, the model and inference remain equally valid for any other partition of multivariate outcomes. The construction is asymmetric, assuming that one type of experimental units is a natural reference for a generative model of co-location.

We suggested the use of PYP priors with negative discount parameter to induce an upper bound on the number of identified clusters. The choice was motivated by the application, based on the resolution of cell subtypes that we are interested in (and that is biologically meaningful to interpret). This use of an upper bound on the number of clusters is not required for the model, and could be dropped if desired, simply by using a non-negative discount parameter.

More complicated structure of the spatial co-localization could be considered if needed. For example, with *J >* 2 sets of experimental units, instead of taking one type of unit as a reference one could allow for co-localization between all possible pairs of clusters of different sets of unit. Formally this could be introduced by using, for example, a CAR type of model across all types of experimental units.

Finally, several limitations remain. One limitation is the assumed a priori independence of cluster locations. For easier interpretation it could be desireable to instead consider a repulsive prior to favor distinct and well separated clusters [38–40]. Another important limitation in the context of the application to the CRC data is the multi-step nature of the bioinformatics pipeline to impute the spatial annotation of the single cell data based on available spatial transcriptomics data. The pipeline ignores the uncertainty in this mapping. Additionally, we model the gene expression features using principal components, which reduces the dimensionality for e”cient inference, but may not capture all the relevant information in the data. One could consider directly modeling the gene expression, but the choice of sampling model and the inference procedure would need to be carefully considered to account for the high dimensionality and potential correlations in the data. Finally, the model casts the problem as one of pairwise co-location. Alternative approaches could construct networks of cell sub-types.

## Appendix

## Appendix A: Proof of propositions in Section 3.2

### Proof of Proposition 3.1

*Proof: Note that the following two models are equivalent*.

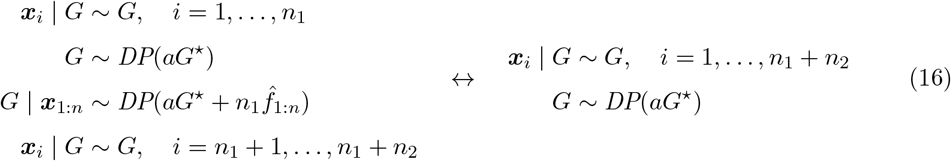

*where* 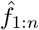 *is the empirical distribution of* ***x***_1:*n*_. *When a*_2_ = *a*_1_ = *a, b*_2_ = *n*_1_, *and* 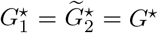, *then model (2), (3) and (4) in the main paper reduces to* (16) *above with* ***x***_*i*_ = ***θ***_1,*i*_,*i* = 1,…, *n*_1_ *and* 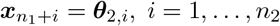.

### Proof of Proposition 3.2

*Proof:* The claim follows from Theorem 3.1 in Guha et al. [31]. For *j* = 1, the SARP model for 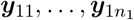 could be written as 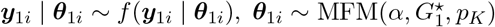, which is a standard MFM model. For *j* = 2, given 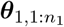 the SARP model for 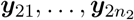 could be written as 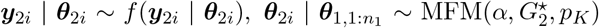, which is also a standard MFM model. Therefore, for the models for both types of units, we can establish the posterior contraction rate following the scheme of proof in [31].

## Appendix B: Details of MCMC algorithm for posterior inference

### Preliminaries for conjugate updates

#### 1. Normal mean with known covariance

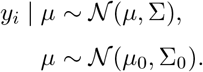

Then

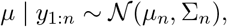

where

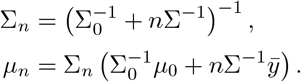

We denote

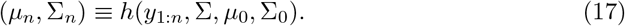

#### 2. Normal–Inverse–Wishart prior

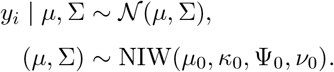

Then

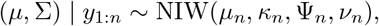

where

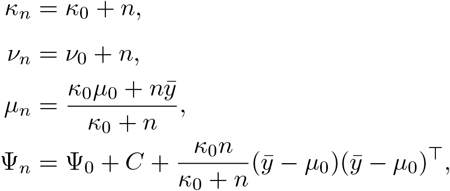

and

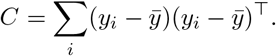

We denote this posterior as

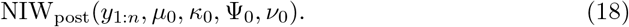

### The SARP model with prior specifications used in simulation and real data analysis

Let *y*_*ji*_ = (*s*_*ji*_, *x*_*ji*_), *ϕ*_*ji*_ = (*m*_*ji*_, *S*_*ji*_), and *ψ*_*ji*_ = (*µ*_*ji*_, ℝ_*ji*_).

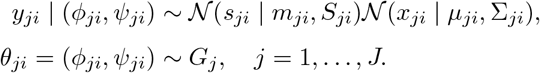

Prior on random measures:

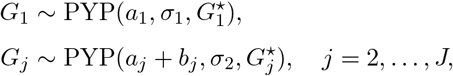

with base measure

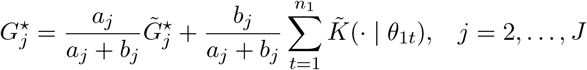

The kernel can be written as:

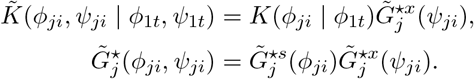

We assume priors:

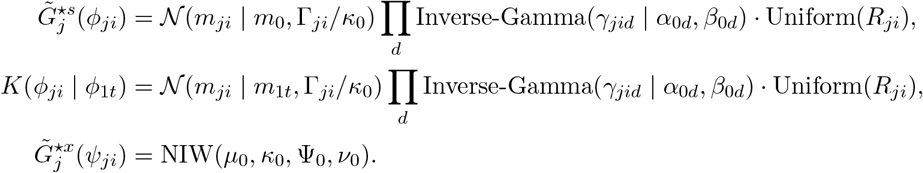

#### MCMC algorithm for posterior inference

Let *L*_*jk*_ = *{*ℓ : *c*_*jℓ*_ = *i, z*_1*i*_ = *k}*

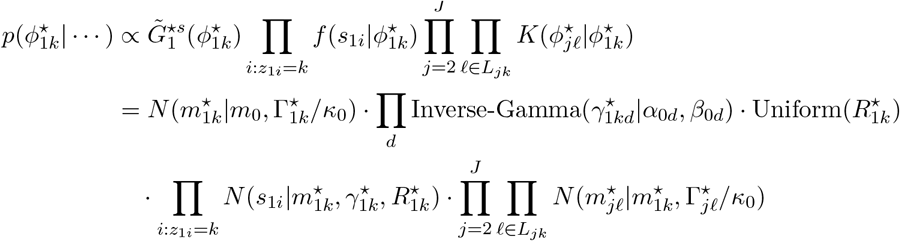

• **1.1 Update** 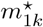

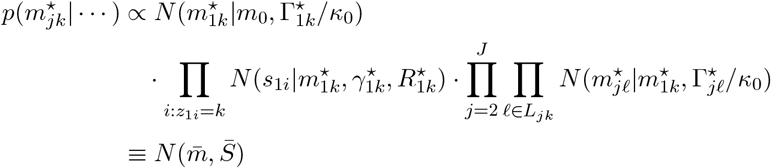

Algorithm for 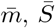:

At a = 0:

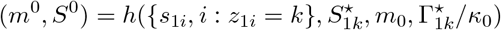

where *h*(·) follows the form in Equation (17).

For *j* = 2, …, *J*:

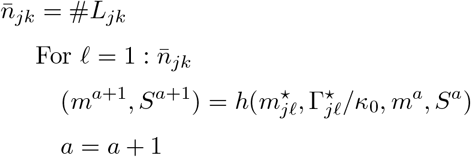

Therefore:

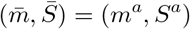

- **1.2 Update** 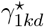

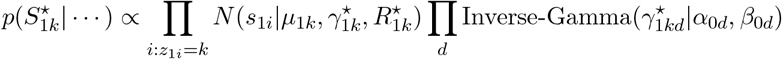

For *d* = 1, …, *D*:

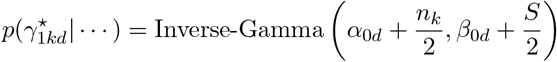

Where

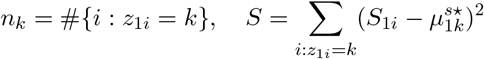

• **1.3 Update** 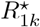

In our 2D case, denote 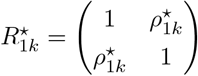

We only need to update univariate parameter 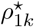.

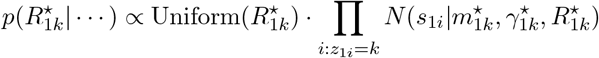

Evaluate conditional posterior probability in the equation above on a grid of values *ρ* ∈ (0, 1) for 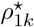 and sample.

• **1.4 Update** 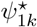

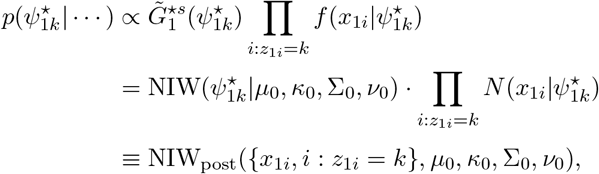

where NIW_post_(·) follows the form in Equation (18).

For parameters 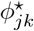, *j* = 2, …, *J, k* = 1, …, *K*_*j*_, we have

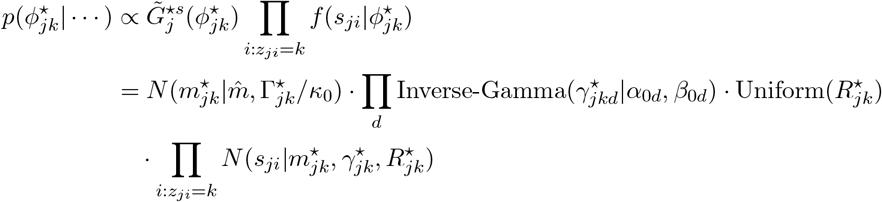

where 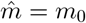 if *c*_*jk*_ = 0, 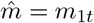if *c*_*jk*_ = *t*.

• **1.5 Update** 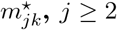

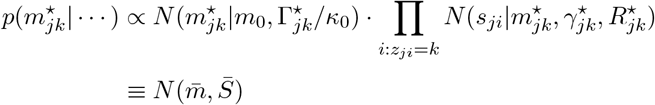

where

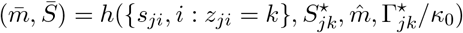

and *h*(·) is given in Equation (17).

• **1.6 Update** 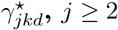

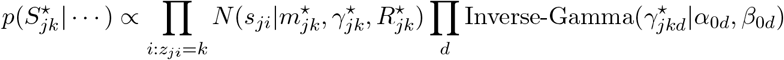

For *d* = 1, …, *D*:

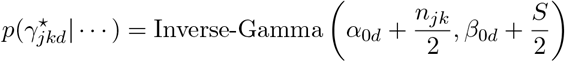

where

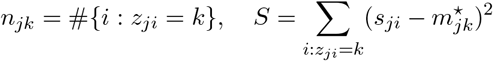

• **1.7 Update** 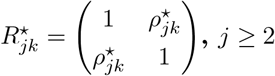

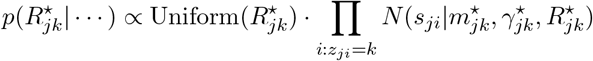

Evaluate conditional posterior probability in the equation above on a grid of values

*ρ* ∈ (0, 1) for 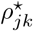 and sample.

• **1.8 Update** 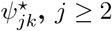

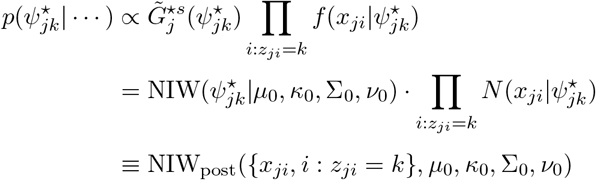

where NIW_post_(·) is given in Equation (18).

• **2. Update** *z*_*ji*_ **for** *i* = 1, …, *n*_1_

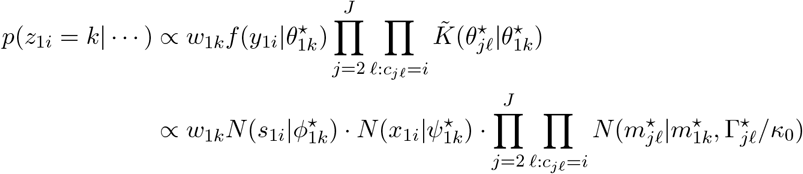

• **3. Update** *z*_*ji*_ **for** *i* = 1, …, *n*_*j*_, *j* = 2, …, *J*

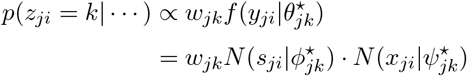

• **4-5: Directly given in section 3.4**

## Appendix C: Details of semi-simulation results

Figure C1 shows the simulation truth and inference results for the semi-simulated data. Inference under the GMM method is implemented and results are shown as benchmark.

**Figure C1:**
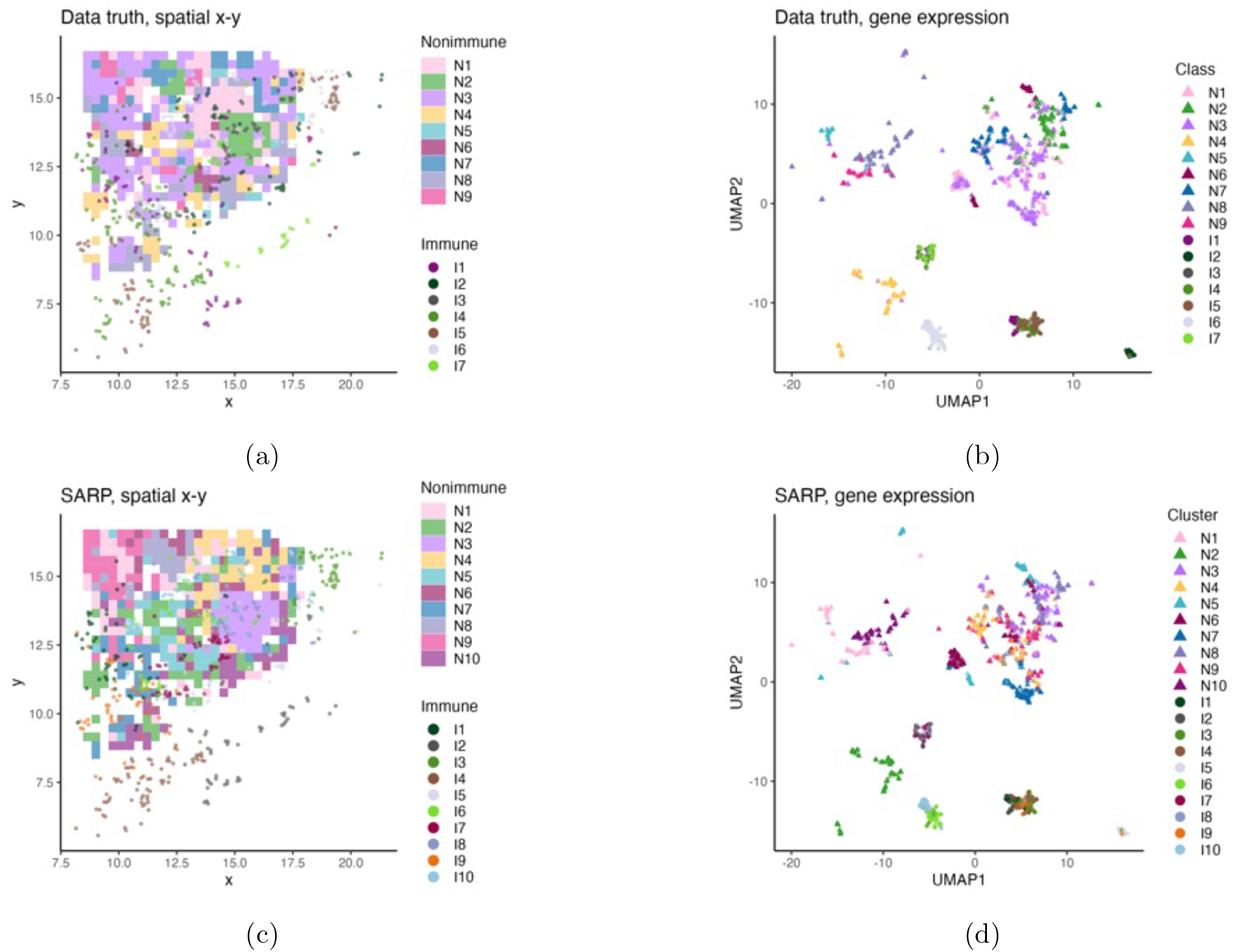
Simulation truth and the estimated partitions under the SARP model for the semi-synthetic data. In the first column the cells are plotted w.r.t. spatial coordinates and in the second column w.r.t. gene expressions. In the first column, grid cells are filled with (background) colors to indicate non-immune cell subtypes, and dots are colored to indicate immune cell clusters. In the right column bullets indicate immune cells, and triangles are non-immune cells, and both are color coded for clusters.

**Figure C2:**
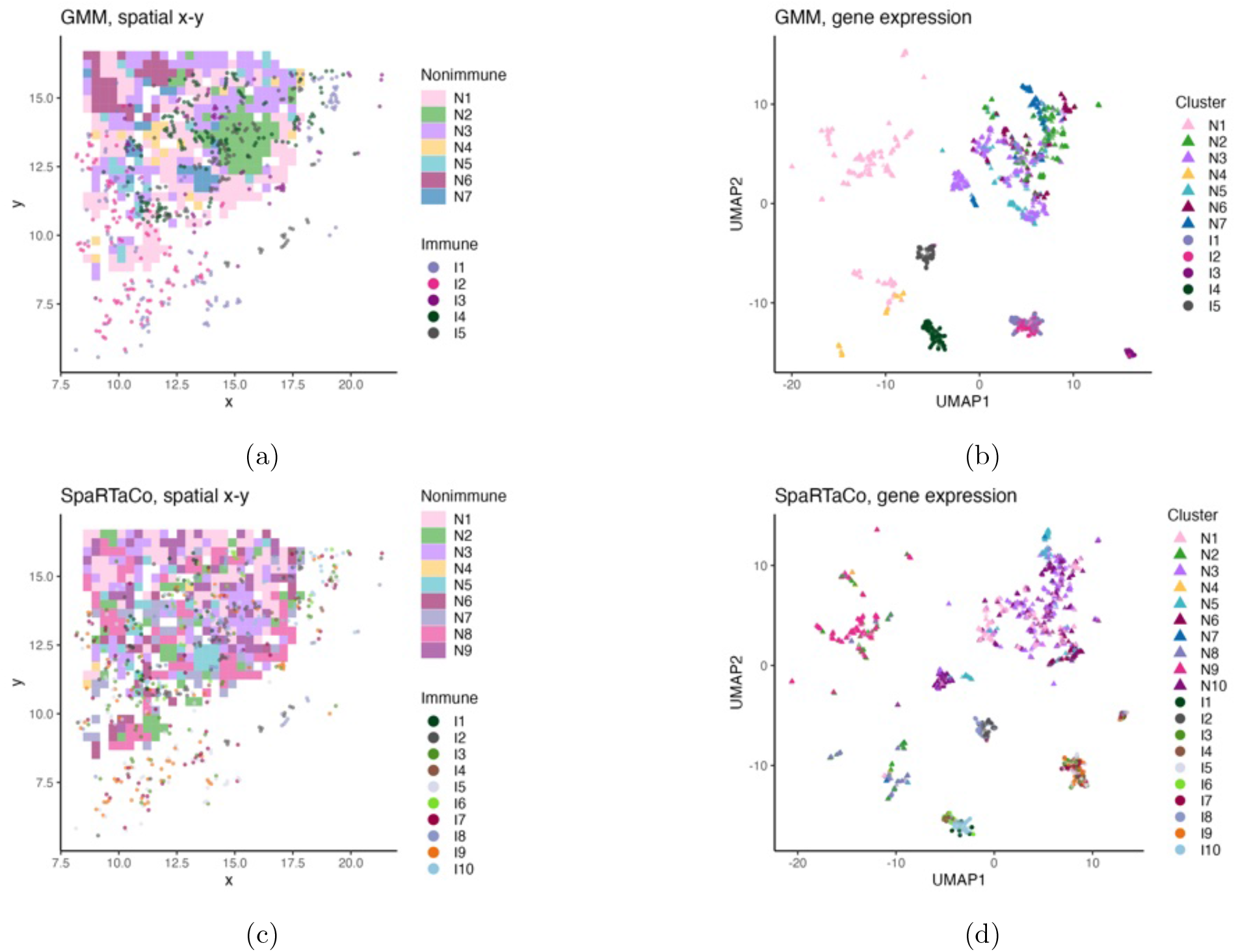
Same as Figure C1(cd) for inference under the GMM (panels (a) and (b)) and under SpaRTaCo (panels (c) and (d)).

## Appendix D: Details of CRC data analysis results

Figure D1 shows spatial alignment for tumor cell subtypes not shown in Figure 5b. For some non-immune subtypes, like Epi2, no immune cell subtypes are found to be spatially aligned.

**Figure D1:**
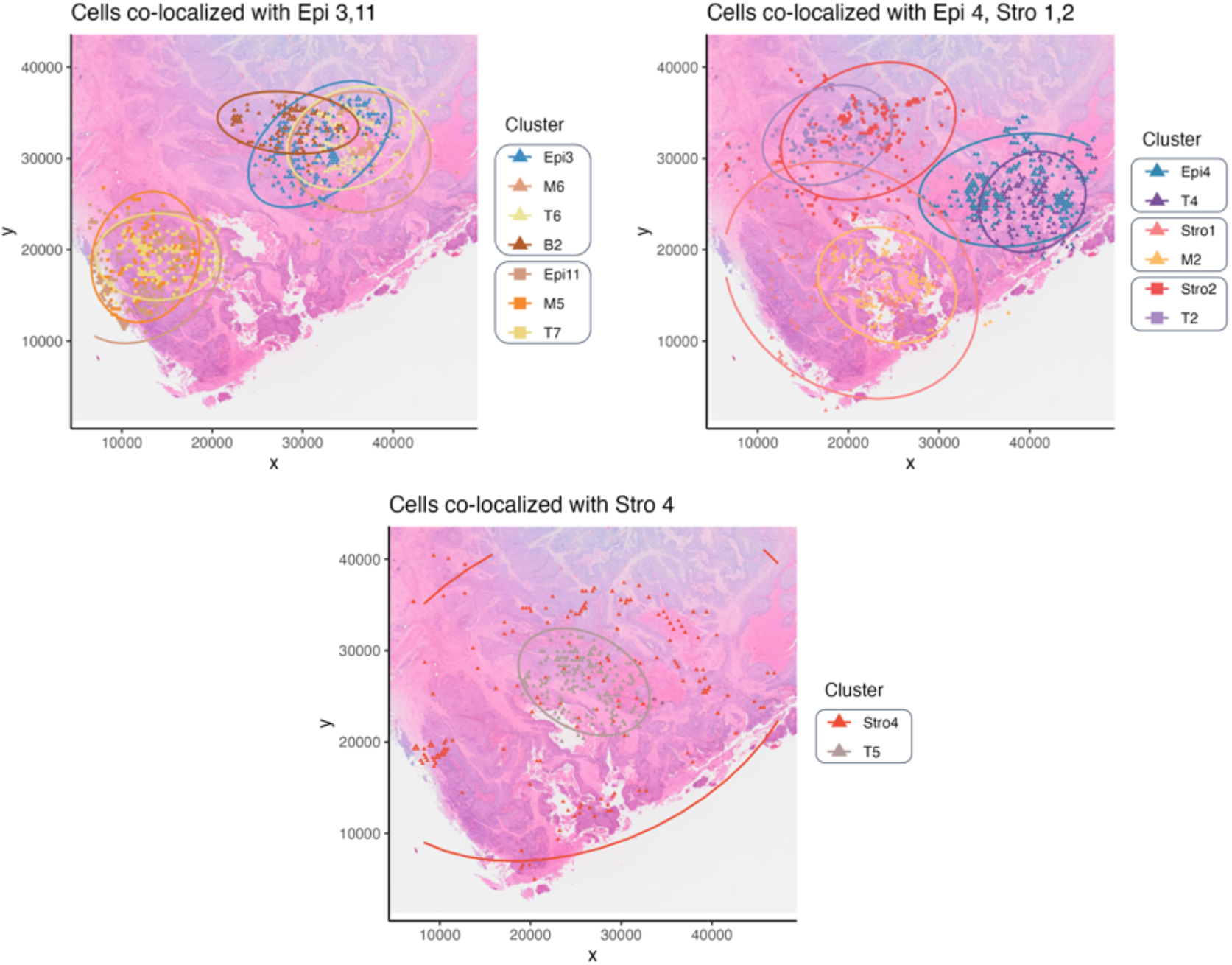
Same as Figure 5b, for immune cells co-localized with non-immune cell clusters that were not shown in Figure 5b.

Figure D2 shows spatial alignment for tumor cell subtypes not shown in Figure 6(b). For some tumor subtypes, like Epi1, no immune or stromal cell subtypes are found to be spatially aligned.

**Figure D2:**
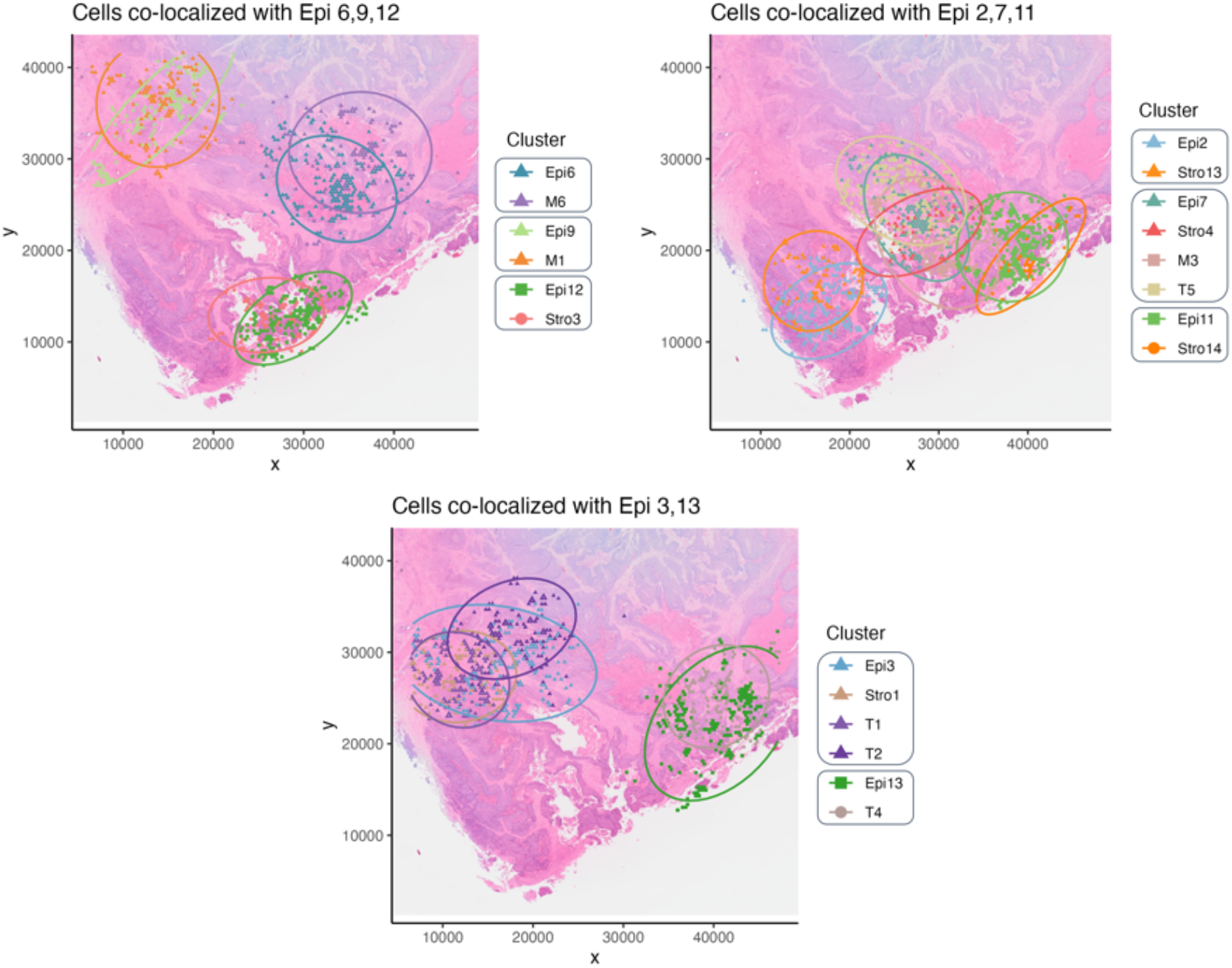
Same as Figure 5b, for immune and stromal cells co-localized with epithelial cell clusters that were not shown in Figure 6b.

Figure D3 shows the posterior probability of immune cells locating independently with tumor cells, and the posterior probability of number of clusters.

**Figure D3:**
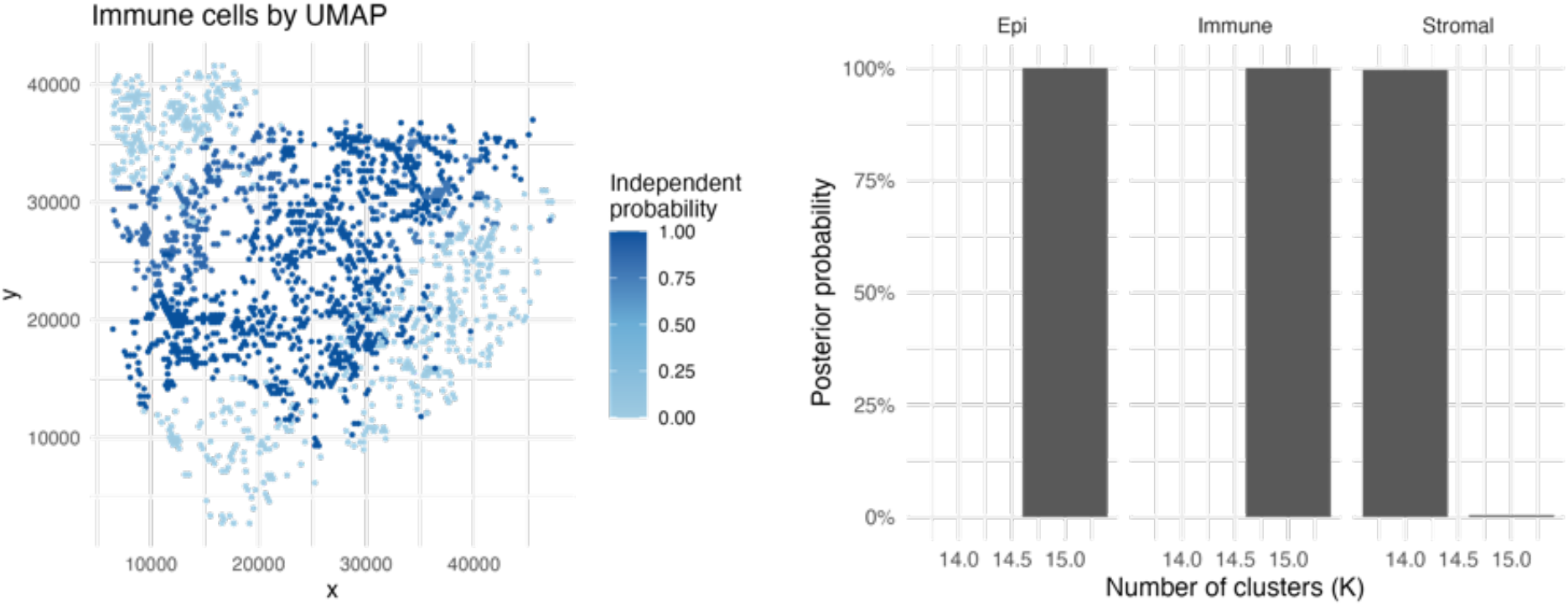
Uncertainty quantification in CRC real data analysis. The left panel shows the posterior probability of the immune cells being spatially independent of tumor cell clusters. Immune cells are plotted with respect to their spatial coordinates. The right panel shows the posterior distribution for the number of clusters for Epi, stromal, and immune cells. All three are degenerate at *K* = 15.

